# Bacterial cell wall biosynthesis is controlled by growth rate dependent modulation of turgor pressure in *E. coli*

**DOI:** 10.1101/2023.08.31.555748

**Authors:** Avik Mukherjee, Yanqing Huang, Seungeun Oh, Carlos Sanchez, Yu-Fang Chang, Xili Liu, Gary Andrew Bradshaw, Nina Catherine Benites, Johan Paulsson, Marc W. Kirschner, Yongjin Sung, Jens Elgeti, Markus Basan

## Abstract

The cell wall is an essential cellular component of bacteria and the target of many antibiotics. However, how bacteria regulate the rate of cell wall biosynthesis as growth rates change remains unresolved. In *E. coli*, cell wall growth was thought to proceed independently from turgor pressure^1^, the osmotic pressure that the cytoplasm exerts on the cell wall. Here, we uncover a striking increase of turgor pressure with growth rate. Modulating turgor pressure and measuring cell wall biosynthesis, we find that turgor pressure is directly controls the rate of cell wall biosynthesis. The picture that emerges is that turgor pressure is largely generated by counterions of negatively charged cellular biomass. The increase in turgor pressure with growth rates results from more ribosomes and therefore higher concentrations of negatively charged ribosomal RNA. Elegantly, the coupling between biomass composition, turgor pressure and cell wall biosynthesis simultaneously explains how bacteria achieve homeostasis of cytoplasmic crowding and how they regulate the rate of cell wall biosynthesis across growth rates.

---

The bacterial cell envelope is one of the most complex chemical structures in nature and its synthesis and expansion involves myriad biochemical processes. Mis-regulation of either cell wall remodeling or cell wall precursor biosynthesis quickly leads to lysis and cell death, a vulnerability that is exploited by some of our most important classes of antibiotics. But how bacteria control the rate of cell wall biosynthesis and expansion across changing growth rates and conditions remains unclear. A fine-tuned regulatory coordination between cell wall growth and biomass production rates is thought to exist, but molecular interactions and molecular players that could mediate this coupling have not been discovered^3,5^. Turgor pressure, on the other hand, is the osmotic pressure exerted by the cytoplasm on the cell wall and an essential property of all prokaryotic cells; it is reasonable to think that it plays a role in driving cell wall expansion. However, experiments in *E. coli* applying periodic osmotic shocks to growing bacteria found that cell volume very quickly recovered after cells exited plasmolysis^1^, a state with vanishing turgor pressure. From this observation, it was concluded that cell wall biosynthesis continues during plasmolysis and therefore that turgor pressure plays no role in cell wall expansion in *E. coli* ^1^. Moreover, it is unknown how cells regulate turgor pressure and whether turgor pressure is constant or changes with growth rates. Therefore, it is unclear how bacteria regulate their cytoplasmic density of protein machinery and other biomolecules, in the absence of a coupling between cell wall expansion rate and cytoplasmic biomass concentration. We therefore hypothesized that such feedback mechanisms must exist and sought to identify the interactions between these important physiological parameters.

## Turgor pressure scales with growth rate

To better understand the physiological role of turgor pressure, we decided to quantify the growth rate dependence of turgor pressure that had never been measured previously. Measuring turgor pressure is challenging because pressure exerted by the cytoplasm is directly balanced by the cell wall^2^ and therefore turgor pressure is not easily accessible. One method to assess the magnitude of turgor pressure involves applying osmotic shocks to bacteria to induce plasmolysis. When bacteria are in plasmolysis, the cytoplasm is compressed and detached from the cell wall at the bacterial poles (Fig. 1a). During plasmolysis, cytoplasmic osmotic pressure is directly balanced by external osmolarity. As we have recently shown, the pressure-volume relationship of cells approximately follows an ideal gas law even after equilibration of salt concentrations^3^. Hence, by measuring the fold-change in cytoplasmic volume as a function of the applied osmolarity under such conditions, it is then possible to calculate turgor pressure of the cell before the osmotic shock (Fig. S1a,b). We thus measured the volume change of the cytoplasm (labeled by expression of a fluorescent protein) of hundreds of bacteria in dozens of osmotic shifts to estimate turgor pressure in different conditions (Fig. 1b,c). When we determined turgor pressure of *E. coli* in different growth conditions in this way, we discovered a striking increase of turgor pressure with growth rate (Fig. 1d & Fig. S1).

**Figure 1:**
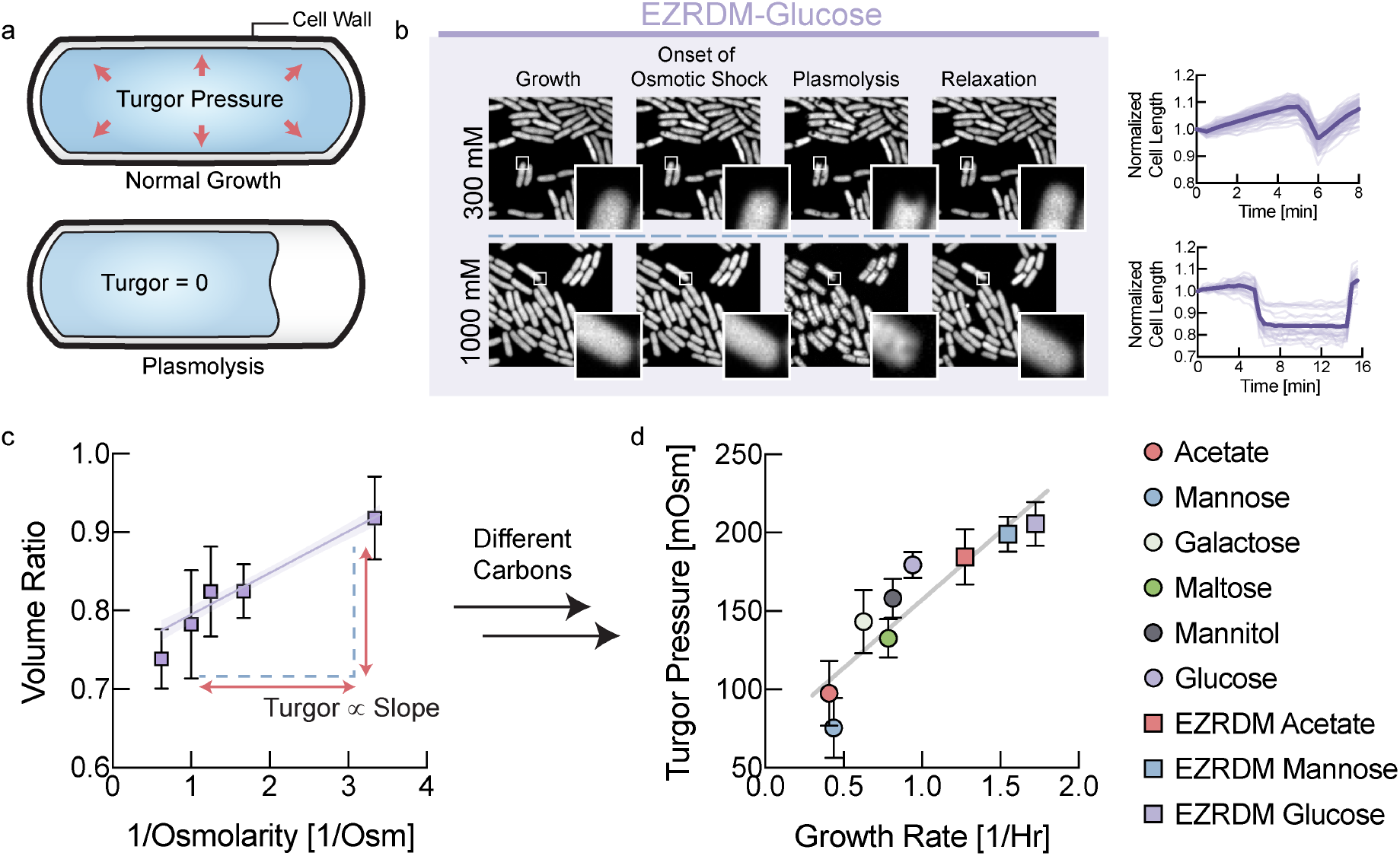
Turgor pressure increases with growth rate. **a**, Osmotic shocks put cells into plasmolysis, a state where the cytoplasm is retracted from the cell wall and osmotic pressure of the cytoplasm is balanced by externally applied osmolarity. **b**, In a microfluidic device, we measured the change in volume of the cytoplasm of hundreds of bacteria after applying osmotic shocks sufficient to induce plasmolysis. **c**, Cytoplasmic volume ratio in the EZRDM glucose condition as a function of inverse of the magnitude of the applied osmotic shock to determine turgor pressure. **d**, Inferred turgor pressure as a function of growth rate. Fits to the Boyle-van’t Hoff equation underlying these data from osmotic shocks of different magnitude are presented in Fig. S1d-l. Error bars represent uncertainty of the fit of the slope to the data in Fig. S1. Growth rates are measured from the same samples that are taken for turgor pressure measurement (See Fig. S1c).

### Turgor pressure controls the rate of cell wall biosynthesis

What is the physiological role of this unexpected coordination of turgor pressure with growth rate? Turgor pressure results in tension in the cell wall and it has been previously established that mechanical forces can lead to plastic deformations of cell shape in *E. coli* ^4^. Therefore, we hypothesized that turgor pressure influences the rate of cell wall biosynthesis. Recently developed D-amino acids become fluorescent only after integration into the cell wall^5^ and therefore enable direct quantification of *de novo* cell wall biosynthesis as a function of time (Fig. 2a). Therefore, we measured the rate of cell wall biosynthesis, quickly after modulating turgor pressure via osmotic shifts in a microfluidic device. Bacteria were pre-grown at steady-state in the presence of regular D-amino acids to prime them for D-amino acid utilization and then shifted to new growth medium that contained fluorescent D-amino acids for cell wall labeling (Fig. 2b). The rate of cell wall biosynthesis in control cells that were shifted to identical osmolarity is shown in Fig. 2c,d.

**Figure 2:**
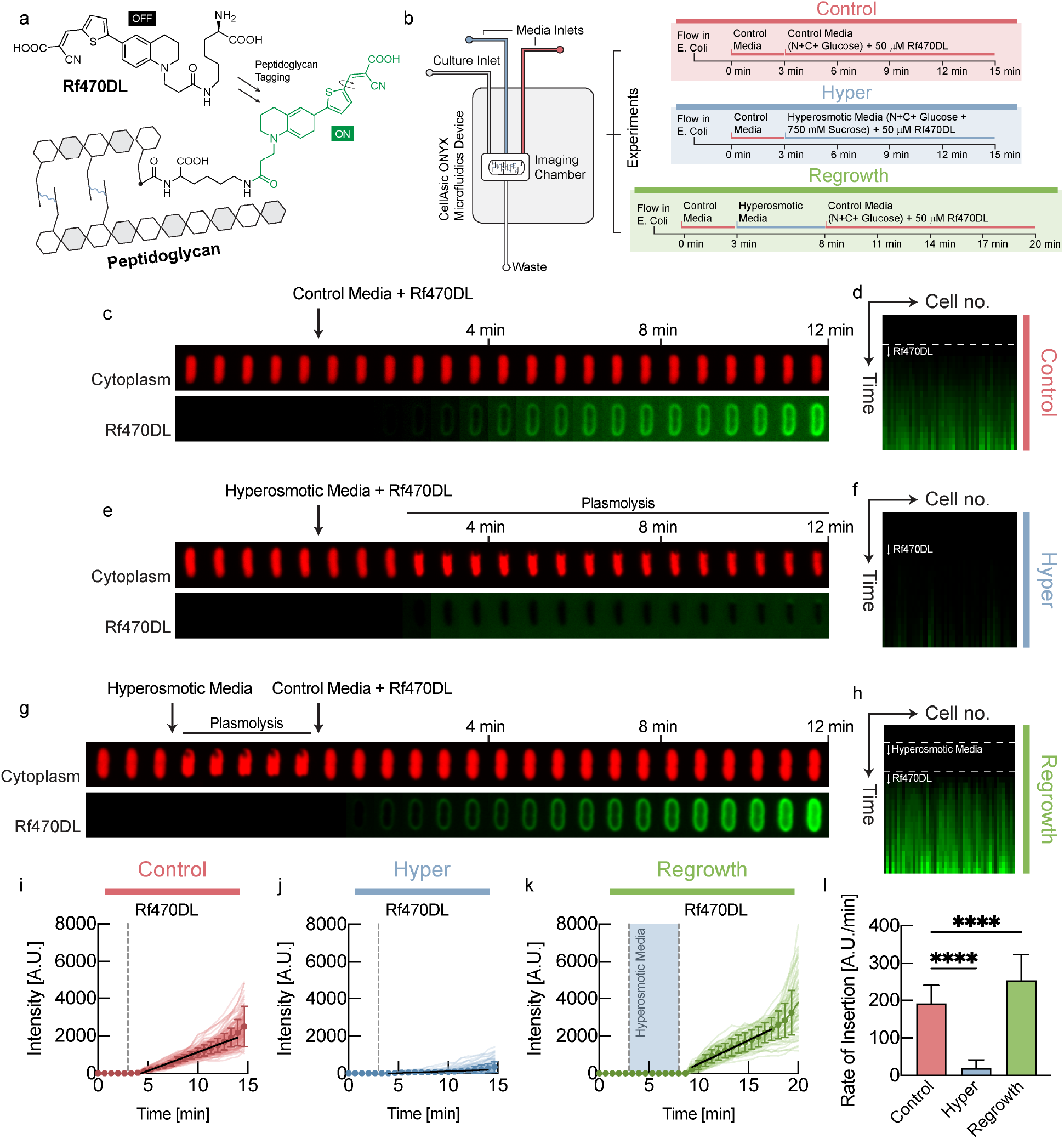
Turgor pressure modulates the rate of cell wall biosynthesis. **a**, Rf470DL ^5^are D-amino acids that become fluorescent after integration into the cell wall^5^. **b**, Using a microfluidic device, we shifted E. coli to medium containing Rf470DL, as well as different media osmolarity. We quantified fluorescence intensity per cell wall length to quantify the rate of cell wall biosynthesis. **c**, An example control cell shifted to identical medium with Rf470DL. **d**, Kymograph of fluorescence intensity of different control cells. **e**, An example cell, shifted to hyperosmotic medium that induced plasmolysis. **f**, Kymograph of fluorescence intensity of different bacyteria in plasmolysis in hyperosmotic medium. **g**, An example cell, shifted back to normal osmolarity after being kept in plasmolysis for 5min. **h**, Kymograph of fluorescence intensity of different bacteria shifted back to normal medium after 5min in plasmolysis. **i**, Fluorescence intensity per cell wall length as a function of time for control cells. **j**, Fluorescence intensity per cell wall length as a function of time for cells in plasmolysis in hyperosmotic medium. **k**, Fluorescence intensity per cell wall length as a function of time for cells shifted back to normal osmolarity after 5min in plasmolysis. **l**, Rate of cell wall biosynthesis quantified from the slope of the traces in i-k.

Strikingly, in cells that were put into plasmolysis by hyperosmotic shock (vanishing turgor), cell wall biosynthesis was virtually undetectable (Fig. 2e,f,j,l). On the other hand, when cells were shifted to hypoosmotic medium (increased turgor pressure), cell wall biosynthesis rate was significantly faster than in control cells (Fig. S4c,e,g,i). Moreover, we also found significantly faster cell wall biosynthesis for cells that were kept in hyperosmotic conditions for several minutes before they were shifted back to normal medium (Fig. 2g,h,k,l). This increased rate of cell wall biosynthesis after return to normal osmolarity is likely caused by increased turgor pressure, resulting from ‘stored’ cytoplasmic biomass, produced during the plasmolysis phase. Indeed, this experiment was also performed by shifting cells to an agar pad for imaging, which confirmed the findings above (Fig. S4), and we also found significantly faster cell wall biosynthesis in shifts to hypoosmotic conditions that increase turgor (Fig. S4d,h,i).

These data suggest that turgor pressure directly modulates the rate of cell wall biosynthesis. Without turgor pressure (plasmolysis) cell wall biosynthesis quickly stopped, while increased turgor pressure (hypoosmotic, shift back from plasmolysis) resulted in faster cell wall biosynthesis. But how does higher turgor pressure interact with the complex biochemistry of the cell wall to result in faster cell wall biosynthesis?

### Model of cell wall fluidization by endopeptidases

One way how mechanical coupling could emerge is from cell wall mechano-endopeptidases that remodel the cell wall in a stress-dependent way^4^. It has been shown theoretically that an elastic material with stress-dependent remodeling behaves like a viscoelastic Maxwell material and can be modeled as a fluid on long timescales^18^. However, even constant activity of cell wall endopeptidases can result in cell wall fluidization, as illustrated in Box 1 (top). Similar models of dislocation-mediated growth of the bacterial cell wall have been proposed^6^. In the illustration in Box 1 (top), peptide bonds cut by cell wall endopeptidases (scissors) result in stress relaxation and expansion flow of cell wall material that is under tension due to turgor pressure. This expansion prevents the transpeptidation repair of these peptide bonds by penicillin-binding proteins (green dots) and creates space for the insertion of new peptidoglycan strands. Because the expansion rate determines the availability of sites for insertion of new cell wall material, cell wall biosynthesis rate is directly proportional to turgor pressure. Metabolic pathways that produce cell wall precursors are known to be product-inhibited. Specifically, UDP-N-acetylglucosamine enolpyruvoyl transferase (MurA) is feedback-inhibited by UDP-MurNAc^7^. When cell wall precursors cannot be integrated into the cell wall (for example due to low turgor pressure), they accumulate in the cytoplasm and shut off their own biosynthesis. Therefore, no fine-tuned regulation of cell wall biosynthesis pathways is required.

Note that this model is consistent with experimental observations that demonstrate that biosynthesis of cell wall precursors is not the limiting factor for cell wall expansion: Neither inhibiting synthesis of cell wall building blocks (fosphomycin) nor preventing their insertion into the cell wall (beta-lactams) stops cell wall expansion and acts bacteriostatic. Instead, both perturbations result in uncontrolled cell wall expansion and lysis, acting bactericidal^8^.

We therefore model the cell wall as a viscoelastic Maxwell fluid which is elastic on short timescales and as a viscous fluid on long timescales along its axis of elongation^9^. As illustrated in Box 1 (bottom), this viscoelastic model naturally results in a volume expansion rate *V̇*/*V* that is directly proportional to turgor pressure.

#### Box 1

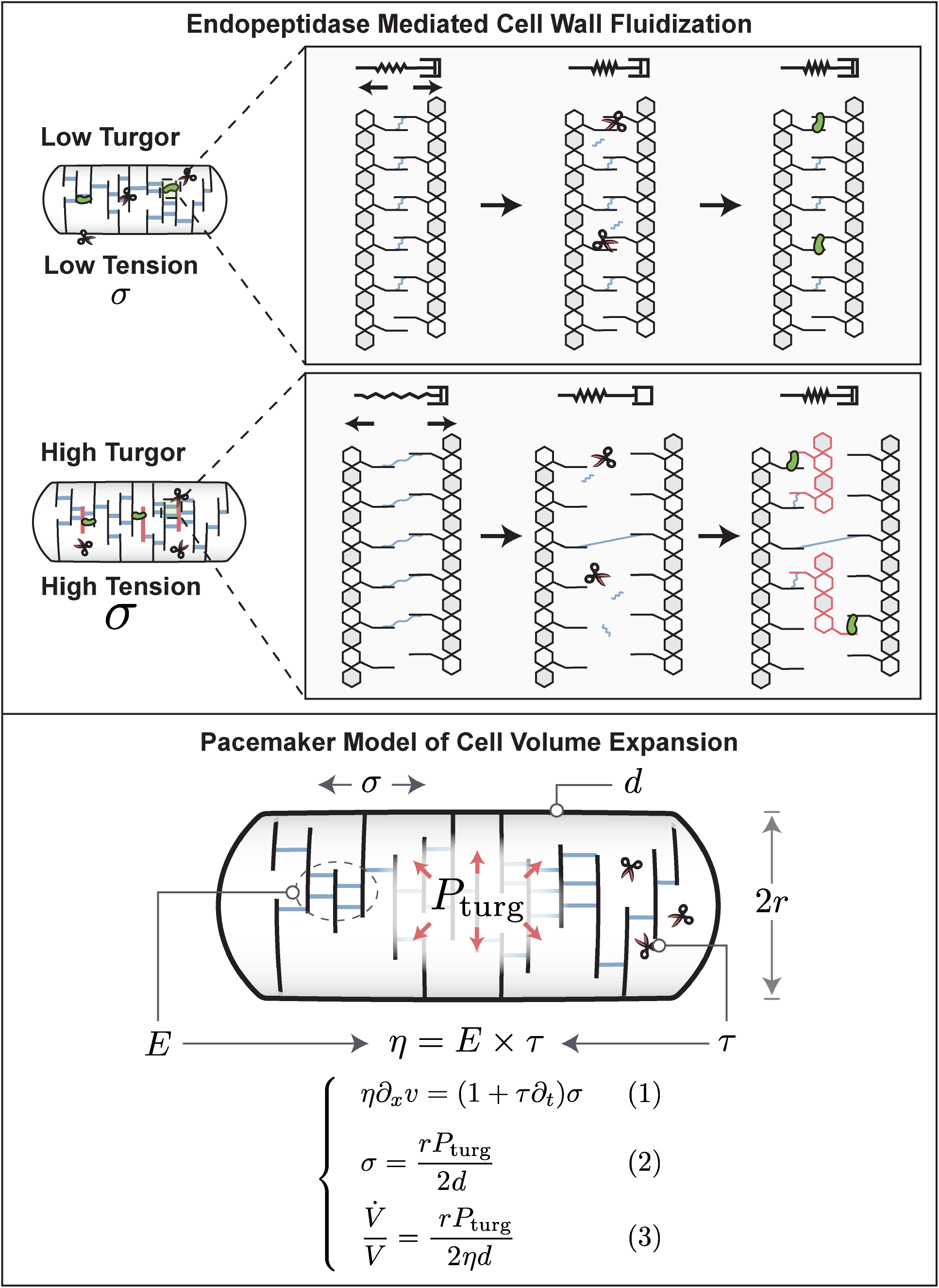

**Endopeptidase mediated cell wall fluidization (top).** Cell wall endopeptidases (scissors) cleave longitudinal peptide bonds that limit cell wall expansion. This leads to expansion of the peptidoglycan network dependent on cell wall stress, generated by turgor pressure (middle panel), enabling insertion of new strands (red, right panel). Endopeptidases thus lead to stress-dependent expansion of cell wall network, resulting in stress relaxation. This constitutes a fluid-like rheology. Therefore, we model the cell wall as a Maxwell viscoelastic material, illustrated by the spring and damper in series. On short timescales such a material behaves like an elastic, whereas on long timescales it exhibits stress-dependent viscous flow that results in cell wall expansion. **Model of cell volume expansion (bottom)**. The cell wall is modeled as a Maxwell viscoelastic material with a constitutive equation, given by Eq. [1]. Turgor pressure P_turg_ from the cytoplasm acts on the cross-section of the cell, generating a tension σ in the cell wall that is given by Eq. [2]. Combining Eq. [1] with Eq. [2] and integrating over the length of the cell, the model predicts a simple expression for volume growth rate, given by Eq. [3]. Model details and derivation of model predictions are presented in Supplementary Note 1, model parameters are summarized in Fig. S5.

### Turgor pressure is required for maintaining biomass density

According to our cell wall model (Box 1, Eq. [3]), the rate of cell wall growth is directly determined by turgor pressure (Fig. 3a). But how is turgor pressure modulated as a function of growth rate (Fig. 3b)? One possibility is that turgor pressure is determined by cytoplasmic biomass density and that cytoplasmic density increases with growth rate. To test this idea, we measured cellular biomass density using computationally enhanced QPM (ceQPM)^10^ (see Fig. S2-S3 & Materials and Methods. See Fig. S6 for comparison and validation of the QPM method with standard methods). However, we found that biomass density was remarkably constant across a large range of growth rates for cells growing on different substrates and in chemostat (Fig. 3c & Fig. S7). By contrast, hyperosmotic growth conditions that have been shown to reduce turgor pressure^11^, resulted in higher biomass density (Fig. 3d). To control turgor pressure more directly, we genetically titrated glutamate. Glutamate is the most abundant intracellular metabolite with intracellular concentrations that should make it a significant contributor to turgor pressure^12^. We used a strain developed by the Hwa lab, in which glutamate producing enzymes were placed under an inducible promoter ^13,14^. Indeed, by titrating intracellular glutamate, it was possible to control cellular biomass density (Fig. 3e). But if biomass density is almost perfectly constant as we observe Fig. 3c, how is growth rate dependent turgor pressure generated and regulated?

**Fig. 3:**
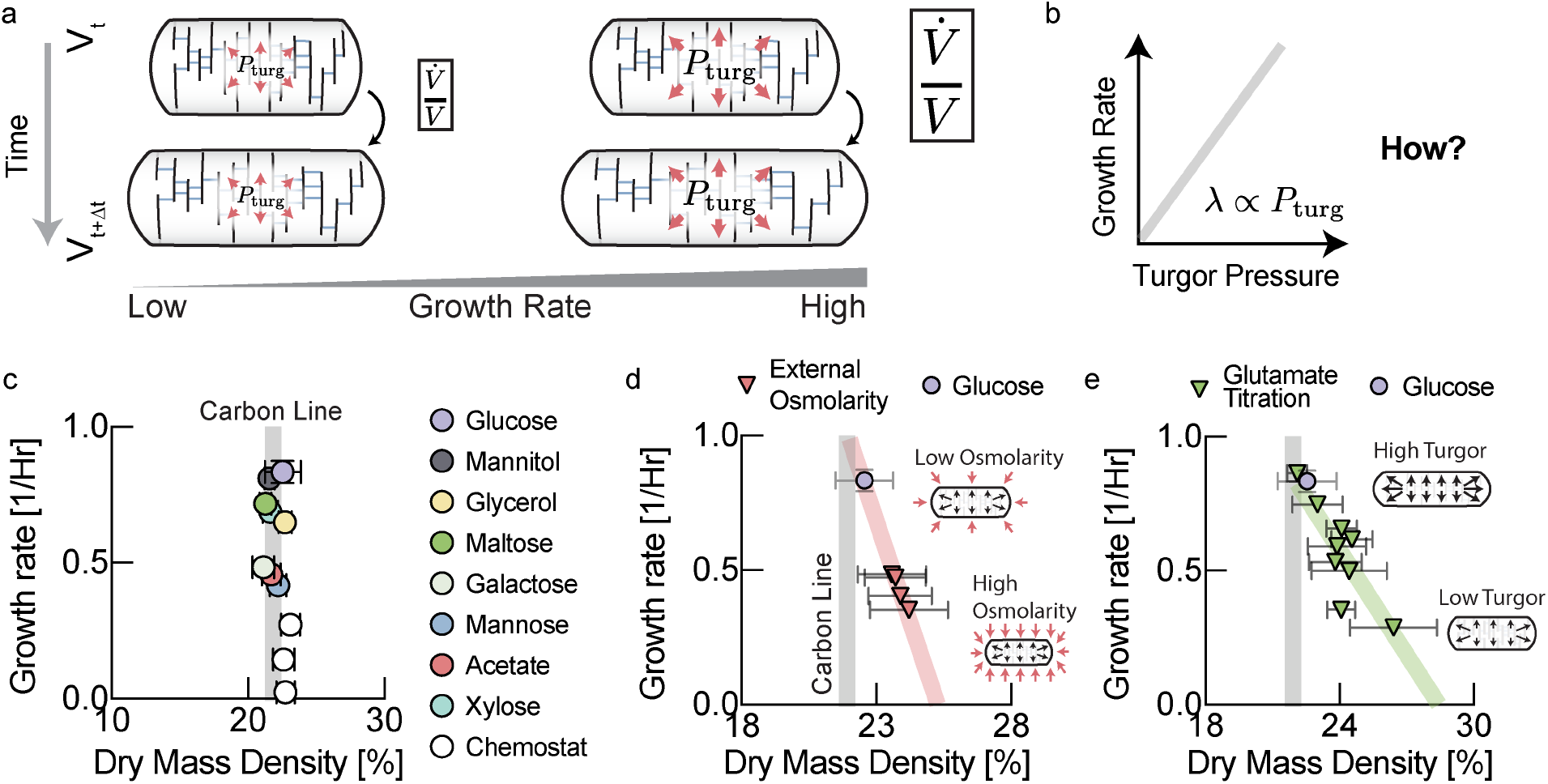
Turgor pressure affects biomass density homeostasis. **a**, We hypothesized that the physiological role of the observed increase in turgor pressure with growth rate in Fig. 1 is to coordinate growth rate of cell volume with growth rate of biomass. **b**, A linear increase of turgor pressure with growth rate ensures that biomass density remains constant across growth rates. **c**, Quantitative phase microscopy measurements show that biomass density is indeed remarkably constant across a wide range of growth conditions. Data for rich defined media is shown in Fig. S7. Growth rates are averages of biological replicates. Error bars in growth rate are the standard deviation from these biological replicates. Plotted dry mass density values result from averaging mean dry mass densities for the measured cell population from different biological replicates. Error bars in dry mass density were determined by propagating the standard deviations of the single cell distributions of the different measurements (Acetate: 3 measurements from n=3 biological replicates; Glucose: 5 measurements from n=4 biological replicates; Glycerol: 6 measurements from n=6 biological replicates; Galactose: 1 measurement from 1 biological replicate; Maltose: 9 measurement from 7 biological replicates; Mannitol: 5 measurements from n=4 biological replicates; Xylose: 3 measurements from n=3 biological replicates; Chemostat conditions: 2 measurements from 2 biological replicates for each data point). Carbon line represents the average dry mass density of different carbon conditions shown in this panel except chemostat experiments **c**, Cells become dense and growth rates slow down when grown in hyperosmotic media that reduce turgor pressure^11^. Individual biological replicates plotted. Error bars are the standard deviation of the single cell distribution of the biological replicate. **d**, Reducing turgor pressure by titration of intracellular glutamate results in higher biomass density and slower growth, similar to the effect of external osmolarity. Individual experiments for different induction levels and biological replicates were binned by their measured growth rate (increments of 0.05/Hr). Error bars were determined by propagating the standard deviation of the single cell distributions from the individual experiments in each bin.

### Increased turgor pressure results from increased intracellular potassium concentrations

All cytoplasmic solutes contribute to turgor pressure, but their individual contribution is approximately given by their cytoplasmic molar concentration. Therefore, we decided to focus on highly abundant osmolytes, based on previous findings^11,15^ (Fig. 4a). The most abundant cellular metabolite is thought to be glutamate. However, intracellular glutamate concentrations have been measured by many different groups and appear to be roughly constant, as we observe (Fig. S8l), or decrease with increasing growth rates under carbon limitation^16^. Therefore, glutamate cannot account for the observed increase of turgor pressure with growth rate. Potassium on the other hand is by far the most abundant cellular osmolyte in *E. coli* ^11^. To test if the observed increase in turgor pressure originated from changes in potassium concentrations, we measured intracellular potassium for *E. coli* growing at different growth rates using mass spectrometry(ICP-MS). Indeed, we observed a significant fold-change in intracellular potassium concentration with growth rate (Fig. 4b & Fig. S8a-i) that can account for the observed changes in turgor pressure (Fig. 1d). Note that absolute potassium concentrations that we measure (see Fig. S8i) are most likely a lower bound of intracellular potassium. This is because cells must be washed with osmo-balanced potassium free medium to remove residual potassium from the filter. However, during this unavoidable washing step that takes around 30s, potassium will diffuse out of the cells into the medium.

**Figure 4:**
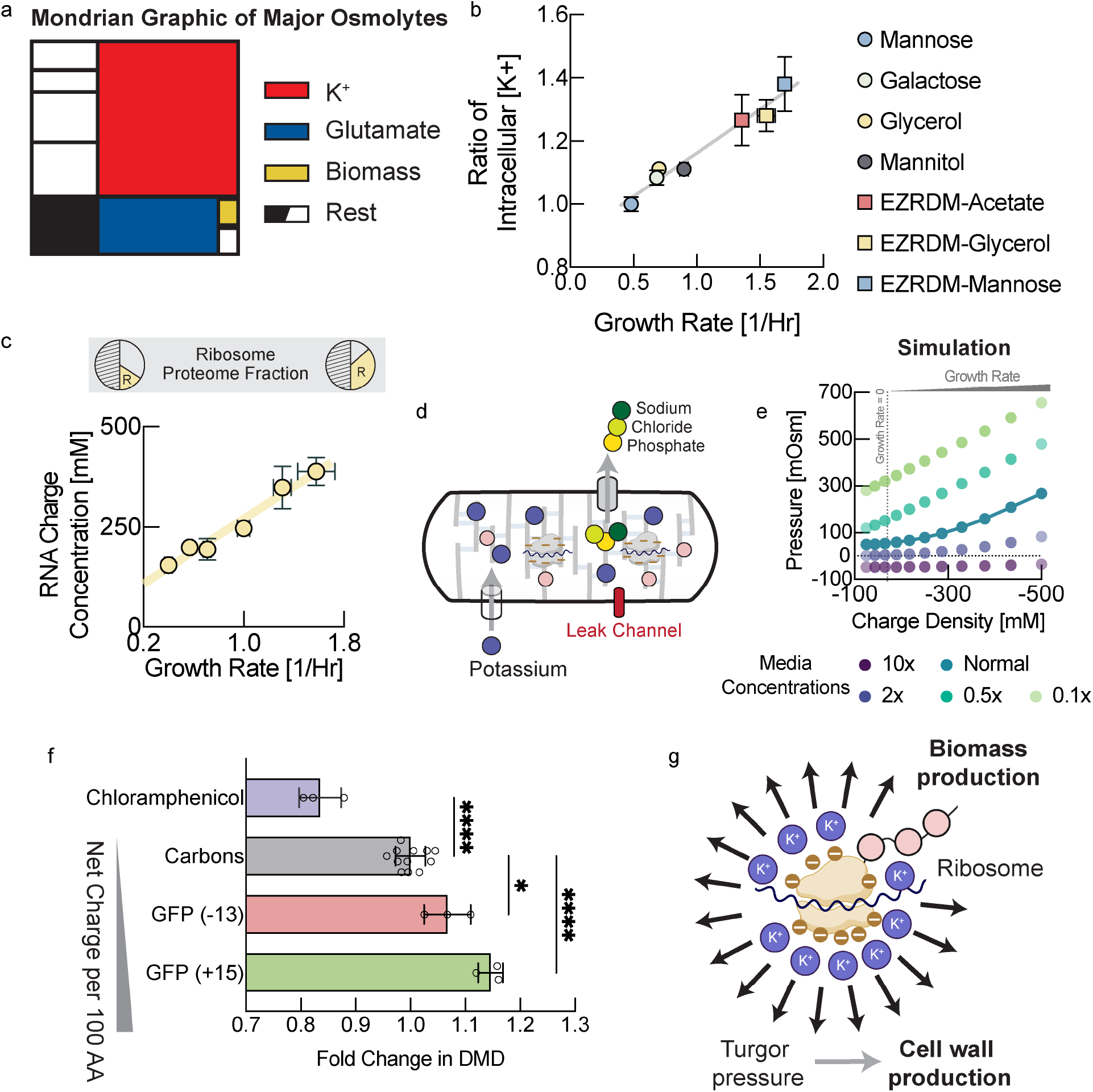
Turgor is modulated by ribosomal counterions to control cell wall expansion. **a**, Schematic illustration of major cellular osmolytes. Potassium and glutamate contribute most to osmotic pressure, whereas the direct contribution from biomass is relatively minor. **b**, Intracellular potassium measured using inductively coupled plasma mass spectrometry (ICP-MS) in low potassium medium in different growth condition. Each data point is determined from several measurements along the growth curve (see Fig. S8 for individual measurements, two biological replicates for each condition). Note that potassium diffuses out of the cell during the mandatory washing step with potassium free medium. Therefore, we plot only relative changes of intracellular potassium across conditions. Error bars represent uncertainty of the fit in Fig. S8. **c**, RNA/protein ratio changes with growth rate according to growth laws. We converted RNA/protein measurements, including error bars, from Scott et al. ^20^ to charge concentrations using total protein and total dry mass measurements from Basan et al. ^21^ (see Fig. S8j,k), as well as dry mass density measurements from this work. **d**, Illustration of the mechano-electro-osmotic model adapted from mammalian cells^3^ for bacteria. For simplicity, we assume expression of a constant proteome fraction of potassium importers, a constant proteome fraction of exporters of all other ions, as well as a constant proteome fraction of passive ion channels. **e**, Numerical solution of the mechano-electro-osmotic model. Predicted turgor pressure is plotted against intracellular charge concentration. There is a baseline of negative charge concentration due to the net charge or protein, glutamate and a growth rate independent baseline level of RNA/DNA (see panel c). In addition, with increasing growth rate there is a linear increase of RNA from ribosomal RNA, as shown in panel c (and DNA from multiple replication forks). Hence, the charge concentration is a proxy from growth rate. **f**, Biomass density increases continuously with overexpression of more positively charged proteins demonstrating the central role of charge balance for biomass density homeostasis. RNA carries one negative charge per nucleotide, whereas proteins are much less negatively charged. Biomass density increases with protein overexpression and the magnitude of this increase depends on net protein charge. Conversely, sublethal doses of chloramphenicol result in higher concentrations of ribosomal RNA^19^ and cause lower biomass density. Because these perturbations result in very long cells of irregular width, biomass densities were determined using Threshold Iterative Volume (TIV) method (see Fig. S9a and Materials and Methods). Biological replicates shown as individual data points. Error bars are standard deviations from these biological replicates. P-values: **: 0.0042; ****: <0.0001. **f**, Ribosomes constitute the central hub of growth control of biomass and the cell wall. They control biomass production rate but also cell wall expansion rate via turgor pressure generated by their counterions.

### Turgor pressure is generated by biomass counterions and modulated by ribosomal RNA

But how is the concentration of intracellular potassium regulated and what causes this increase in intracellular potassium in coordination with growth rate? Potassium is positively charged, while the net charge of the cytoplasm must be neutral. One way to explain charge neutrality is that lipid membranes have a very small electrical capacitance, so a tiny misbalance of net charge results in a large membrane potential that will in turn attract ions, balancing the net charge. Although there is a tiny misbalance of cytoplasmic charge that results in membrane potential, charge neutrality can be assumed for all practical purposes (a standard assumption in electrophysiology^17,18^). Hence, growth-rate dependent potassium concentrations must be balanced by an equivalent negative growth-rate dependent charge concentration in the cytoplasm. The concentration of the major intracellular anion glutamate has been shown to be roughly constant or decreasing with growth rate under carbon limitation^16^. Moreover, glutamate is not sufficiently abundant to balance potassium concentrations (see Fig. S8i,l). According to studies of cellular osmolytes ^11,15^, there are no other negatively charged salt ions or metabolites that are thought to be sufficiently abundant to balance the charge concentration of potassium.

On the other hand, cellular biomass including protein, DNA and RNA carries a net negative charge at physiological pH. According to our estimates, the biggest negative contribution to cytoplasmic charge comes from RNA, as each nucleotide carries one negative elementary charge. Moreover, cellular RNA content as a fraction of biomass is well-known to increase with increasing growth rates^19^. This is the result of an increase in ribosomal RNA, which makes up the lion’s share of cellular RNA. According to bacterial growth laws, higher concentrations or ribosomes are required to support faster growth rates. Thus, we converted the RNA to protein ratios across growth rates, measured by Scott et al. ^20^, to an intracellular charge concentration by using our own measurements of biomass density in this work, combined with earlier measurements of total protein and total dry mass across growth conditions^21^(Fig. 4c). Indeed, the increase in charge concentration with growth rate from this analysis was very large (>100mM). This charge must be balanced by positively charged counterions (potassium) for charge neutrality of the cytoplasm and indeed the measured increase in intracellular potassium concentration has a comparable magnitude (Fig. S8i). We also estimated the net charge of ribosomal proteins and confirmed that their net positive charge compensates only about 10% of the net charge from ribosomal RNA (Table S1). Moreover, biophysical experiments and theory suggest that at most 50% of counterions are condensed on charged macromolecules like RNA and DNA and thus at least 50% are osmotically active and contribute to turgor pressure^22–24^.

### Machano-electro-osmotic model of turgor pressure

Using a mechano-electro-osmotic biophysical model of the cell that we developed for mammalian cells^25^, we wanted to test if bacteria could achieve scaling of turgor pressure proportional to growth rate via changes in their ribosomal concentration. In brief, this model is based on first physical principles: We solve the Nerst-Planck equation for every ion species together with considering active ion transport. Then, combined with charge balance and mechanical force balance, the model predicts intracellular ion concentrations, membrane potential and turgor pressure. We made the simple assumption active import of potassium and active export of all other inorganic ions (Fig. 4d), in order to qualitatively match experimentally observed ion concentration^11,15^. The only other assumption of the model is a constant expression ratio of ion pumps and channels (e.g. due to constant proteome fractions).

Indeed, using this model, we found a large parameter regime where substantial turgor pressure is generated by ribosomal RNA. Turgor pressure correctly scales with increasing growth rates (increasing charge concentration due to the proteome fraction of ribosomal RNA) (Fig. 4e). In addition, the model demonstrates that bacteria can generate sufficient turgor pressure to drive cell wall expansion over a large range of media osmolarities (Fig. 4e, different colors). Consistent with experimental observations (Fig. 3d), changes in media osmolarity result in changes in turgor pressure. These changes in turgor must be compensated by changes in cytoplasmic biomass density, according to our cell wall model (Box 1, Eq. [3]). Hence, despite the existence of osmo-adaptive programs in *E. coli*, this simple model is consistent with phenotypes of bacteria at higher and lower medium osmolarities (Fig. 3d & Fig. S9b). Together, these data suggest that rather than being directly sensed and tightly controlled, turgor pressure is an emergent property that is modulated by environmental factors like medium composition.

To test this hypothesis experimentally, we asked how changes in cytoplasmic net charge are reflected in turgor pressure and cellular biomass density. If charge balance were indeed the key determinant of turgor driving cell wall expansion, then it should be possible to modulate biomass density by affecting net charge of biomass. Overexpressing large quantities of proteins of different net charge should therefore differentially affect cellular biomass density. More positively charged protein overexpression should result in a larger increase in biomass density. Using an established system for expression of large quantities of useless proteins ^26^, we expressed positively and negatively supercharged versions of GFP, developed by the Liu lab ^27^. Indeed, we found that biomass density increased with the net positive charge per amino acid of the overexpressed protein (Fig. 4f), just as expected from the hypothesis that turgor pressure is controlled via cytoplasmic charge balance. Conversely, we used sublethal doses of chloramphenicol, which have been shown to result in upregulation of ribosomes and therefore a higher fraction of ribosomal RNA^20^. A shift of biomass composition from protein to RNA should result in an increased negative net charge of cellular biomass and therefore a decrease in biomass density. Indeed, this is what we observed experimentally when we measured biomass density in these cells (Fig. 4f).

### Biomass density homeostasis and regulation of cell wall biosynthesis rate via ribosome-mediated modulation of turgor pressure

By controlling turgor pressure through ribosomal RNA, bacteria elegantly achieve homeostatic feedback control of biomass density and at the same time coordinate cell wall expansion with growth rate: Because turgor pressure is proportional to biomass density (Box 2, Eq. (4)), cells that are too dense have increased turgor pressure. This results in faster volume expansion according to Box1, Eq. (3) and therefore constitutes a negative homeostatic feedback loop to control cellular biomass density (Box 2, top).

On the other hand, bacterial growth laws require ribosome content to scale proportional to growth rates to support efficient self-replication. This increase of ribosomal RNA automatically results in an increase in turgor pressure due to a larger biomass fraction of ribosomal RNA that is also proportional to growth rate (Box 2, bottom & Eq. (4)). Higher turgor then results in faster cell wall expansion according to Box1, Eq. (3), matching biomass growth rate in the cytoplasm. This ensures constant biomass density across growth rates (Fig. 3b). Thus, using this mechanism to control turgor, bacteria achieve both homeostatic feedback control of biomass density, as well as constancy of biomass density across a large range of growth rates, without the need for fine-tuned regulatory coordination between cell wall synthesis and biomass production rates.

#### Box 2

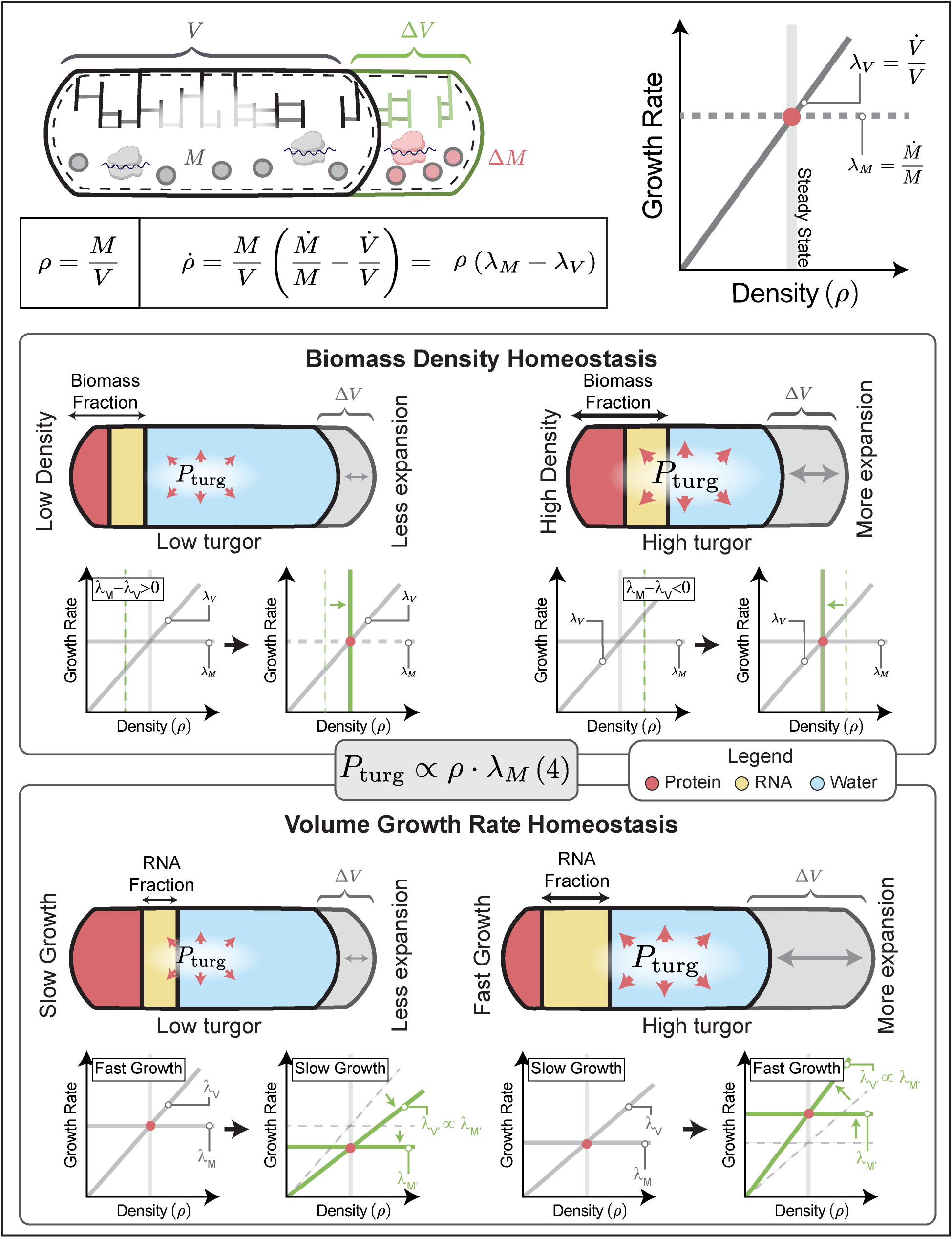

**Electro-osmotic model of ribosome-generated turgor pressure and homeostasis. Top**, Biomass density is determined by biomass growth rate and volume growth rates that must match in steady-state conditions. Because turgor pressure is proportional to biomass density, volume growth rate is also proportional to biomass density, according to Box 1, Eq.[3]. **Middle:** Illustration of dilute cells (left) and dense cells (right). Turgor pressure generated by counterions biomass results in turgor pressure proportional to biomass density (Eq. (4)). Higher biomass density then results in higher turgor pressure (Eq. (4)) and thereby faster cell volume expansion (Box 1, Eq. (3)). This creates a homeostatic feedback loop for control of biomass density. **Bottom**, Illustration of slow-growing cells (left) and fast-growing cells (right). Due to growth laws^19^, a larger fraction of biomass is made up of ribosomes in fast-growing cells as compared to slow-growing cells (higher RNA/protein ratio). This results in a higher molarity of counterions per biomass. At constant total biomass density, faster growing cells have a higher counterion concentration and correspondingly a higher turgor pressure (Eq. (4)), as observed experimentally (Fig. 1c). Higher turgor thus results in faster cell volume expansion (Box 1, Eq. (3)), as required for maintaining constant biomass density across growth rates (see Fig. 3c & Eq. (S5)).

### Perturbations of cell wall properties and model predictions

Our model makes a set of predictions about the relationship between important cellular properties. In our model, cell wall rheology is determined by an effective viscosity, given by η = *E τ*, where *E* is the elastic modulus of the elastic network of the cell wall and *τ* is the viscoelastic relaxation time, which reflects cell wall remodeling by endopeptidases. Therefore, one way to affect cell wall viscosity is changing the rate of remodeling of the cell wall due to endopeptidase limitation^28^. As illustrated in Fig. 5a, downregulating the abundance of cell wall endopeptidases should result in higher cell wall viscosity, slowing down volume growth rate. Higher viscosity from lower hydrolase abundance must thus be compensated by a combination of slower growth rates, greater cell widths or higher biomass densities (Fig. 5a, middle).

**Figure 5:**
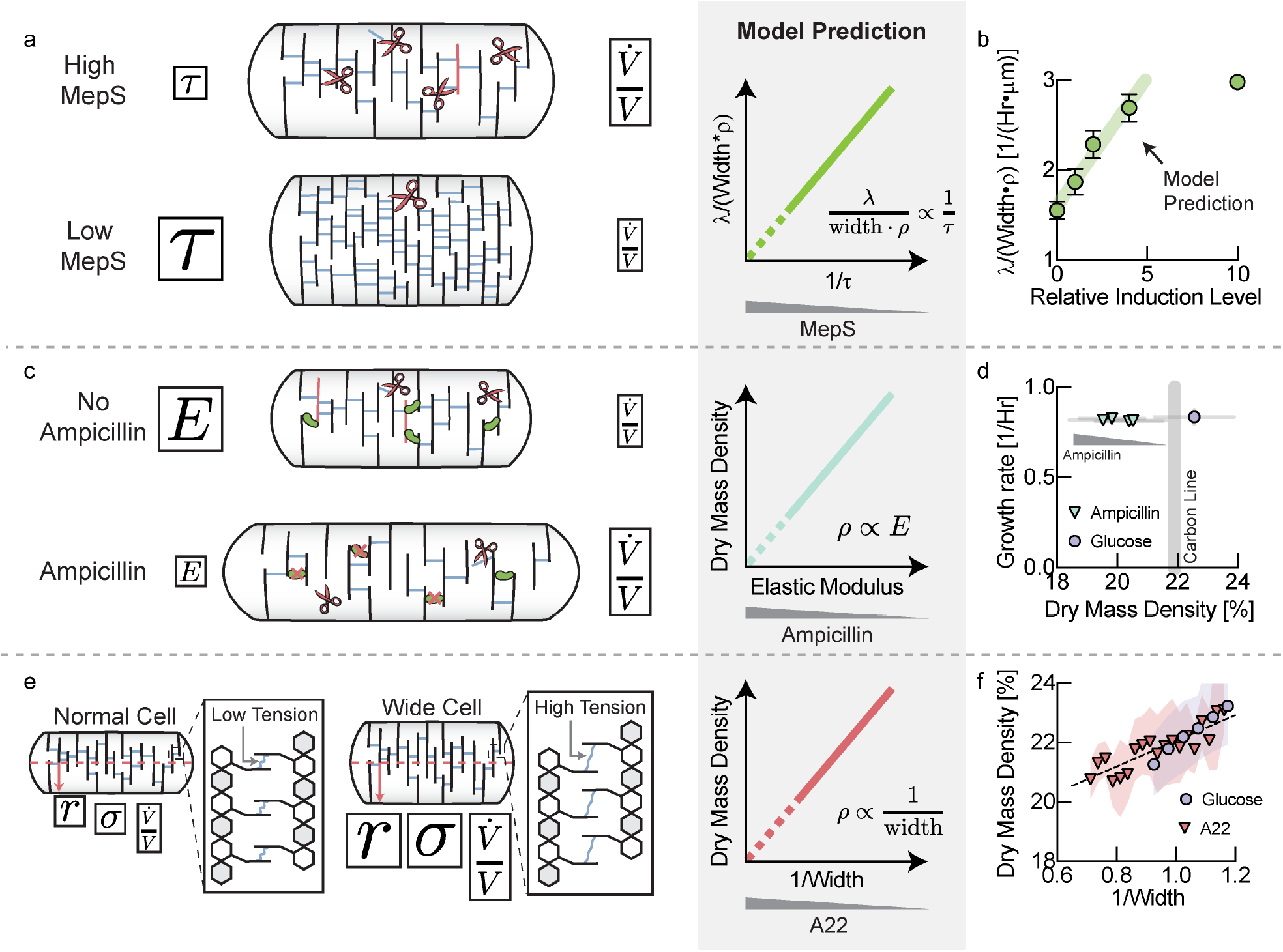
Perturbation of cell wall properties. **a**, Titration of the abundance of cell wall endopeptidases affects the effective viscosity of the cell wall, resulting in a slower volume growth rate with lower expression levels. The model predicts that increasing cell wall viscosity must be compensated by a combination of growth rate, biomass density and cell width (see Supplementary Note 1, Eq. [S13]). **b**, Comparison of experimental data to model prediction. A non-zero y-intercept may emerge from the activity of other cell wall endopeptidases or leaky expression. Error bars are standard deviations of the values calculated from the mean of individual biological replicates. Biological replicates identical to panel C. **c**, Ampicillin affects the crosslinking density of the peptidoglycan network and thereby its elastic modulus. The elastic modulus of the cell wall is proportional to its effective viscosity and therefore controls volume growth rate. The model predicts a direct proportionality between biomass density and the elastic modulus of the cell (see Supplementary Note 1, Eq. [S10]). **d**, Sublethal doses of ampicillin resulted in decreasing biomass densities, while growth rate was unaffected. Error bars represent standard deviation of biomass density from the distribution cells measured in each experiment. **e**, A larger cell width results in higher cell wall tension and therefore a higher volume expansion rate. The model predicts that faster volume expansion rate due to higher cell width must be compensated by a drop in biomass density (see Supplementary Note 1, Eq. [S8]). **f**, Average biomass density of individual cells, binned by their inverse cell width drops with increasing width. The effect is amplified by inhibiting MreB using A22 and also holds when pooling data across different carbon sources (see Fig. S10d). Glucose bin increment: 0.05µm^-1^. A22 bin increment: 0.025µm^-1^. Shaded areas represent the standard deviation of the distribution of cells in each bin.

Recent work identified three cell wall hydrolases that together are essential for cell growth of *E coli* ^29^. We therefore knocked out two of these hydrolases (MepM, MepH) and replaced the chromosomal promoter of the third (MepS) by a linearly inducible expression system^30^. Indeed, we found that at low MepS induction levels resulted a combination of slower growth rates, wider cells and higher biomass densities, consistent with our model prediction (Fig. 5a, left & Fig. S10a-c).

The effective viscosity of the cell wall is not only determined by the rate of remodeling, but also by the elastic modulus of the underlying elastic network (see Box 1). Therefore, affecting the elastic modulus of the cell wall should directly affect cell wall expansion rate and biomass density. Ampicillin is a beta-lactam antibiotic that blocks cell wall crosslinking and insertion of new material into the cell wall^31^. Therefore, cells treated with sublethal doses of ampicillin should have a decreased crosslinking density and a softer, more elastic cell wall (smaller elastic modulus *E* and effective viscosity η, see Fig. 5c). According to Box1, Eq. [3], this decrease in effective viscosity would be expected to be compensated by a drop in turgor pressure and thus, according to Box 2, Eq. [4], drop in biomass density (Fig. 5c & Supplementary Note 1, Eq. [S10]). Indeed, we found that increasing sub-lethal concentrations of ampicillin continuously decreased cellular dry mass density during steady-state growth (Fig. 5d), while growth rates were approximately constant.

Finally, the model also makes a prediction regarding the effect of cell width on biomass density (Fig. 5e). According to Box1, Eq. [3], cell wall expansion rate is directly affected by cell width because turgor pressure creates tension in the cell wall by acting on the cellular cross section. Cell wall tension and the expansion rate of the cell should therefore increase with cell width for a given turgor pressure. This must be compensated by a drop in turgor pressure via a drop in biomass density, according to Box 2, Eq. [4]. Indeed, binning cells in a growing population on glucose minimal medium according to their width, we confirmed that biomass density decreased with cell width (Fig. 5f, blue circles). This single-cell trend was also seen by averaging hundreds of thousands of cells in different growth conditions based on cell width (Fig. S10d). Moreover, when we inhibited MreB, a protein known to be involved in regulation of cell width in *E. coli* ^32,33^, we found a similar, but amplified relationship between biomass density and cell width (Fig. 5f, red triangles & Fig. S10e).

## Discussion

Biomass production and cell wall synthesis are two seemingly disjoint cellular processes that must nevertheless be tightly coordinated. Without such coordination cells would produce either toxic molecular crowding or lyse from a buildup of turgor pressure or thinning of the cell wall network. Our work shows that cell wall expansion is intimately coupled with biomass synthesis in the cytoplasm via turgor pressure. Turgor pressure controls cell wall expansion rate and is generated counterions of biomass, in particular ribosomes, the same machinery that produces cytoplasmic biomass. Ribosomal scaling with growth rate results in an increase in turgor pressure that ensures that cell wall expansion increases with growth rate, allowing cells to grow at constant biomass density. No fine-tuned regulation of cell wall precursor producing metabolic pathways is required in this picture. Instead, simple product inhibition from cell wall building blocks is sufficient to ensure that biosynthetic flux of cell wall building blocks matches requirements from cell wall expansion. Finally, because turgor pressure is generated by biomass counterions, regulation of cell wall expansion by turgor pressure also results in homeostasis of cellular biomass density: Cells that are too dense will expand faster due to high turgor, thus reducing their biomass density. Conversely, cells that are too dilute will expand more slowly, allowing biomass growth to catch up with volume and increasing biomass density.

Interestingly, the role of turgor pressure in coordinating cell wall expansion is consistent with our recent findings that starving bacteria expend the lion’s share of their ATP maintenance budget to maintain plasmolysis and reduce turgor by exporting ions^34^. Plasmolysis is a state with vanishing turgor pressure and according to our model (Box1, Eq. (3)), vanishing turgor is required to stop the expansion of the cell envelope. Hence, even a small buildup of turgor due to the intracellular concentration of macromolecules, metabolites or counterions would result in cell volume expansion in starving bacteria. But in starvation conditions, there is no cell wall precursor synthesis possible, which is required to reinforce the expanding cell wall. Therefore, expansion would quickly result in lysis and cell death. Indeed, this is precisely the phenotype that we observed in starving bacteria after loss of ion homeostasis due to lack of ATP prior to cell death^34^: The cytoplasm of cells that lost membrane potential began to swell, and they exited plasmolysis. We then observed expansion of the cell wall and lysis, accompanied by loss of biomass soon after. Therefore, plasmolysis may be essential to prevent cell wall expansion during starvation.

In conclusion, we uncovered an elegant interplay of basic physiological mechanisms acting in concert to coordinate cell wall biosynthesis rate with biomass production rate in the cytoplasm. While we focused on the model organism *E. coli* in this work, these mechanisms are possibly conserved across evolution, extending to other microbial species^36 37 38–41^. Viscoelastic models of cell wall growth have been successfully applied to plant growth^35^, where the turgor pressure driven cell wall expansion remains the predominant model.

## Acknowledgements

We would like to thank Terry Hwa for many discussions about these topics and much helpful advice and suggestions. We would also like to thank Noah Olsman, Ariel Amir, Joshua Rabinowitz, Jan Skotheim, Becky Ward, Quincey Justman, and Sean Megason for helpful feedback on the manuscript. We would also like to thank Jennifer Waters and Talley Lambert and Anna Payne-Tobin Jost(Nikon Imaging center, Harvard medical school), for valuable comments on image acquisition and processing. We thank Professor Charles Langmuir (Geology department, Harvard University) for giving us access to ICP-MS facility and thanks to Zhongxing Chen for helping us with the ICP-MS experiments. We thank Michael Springer of Harvard Medical School for giving us access to CellAsic microfluidics device and special thanks to Ang Li (Springer lab) for technical troubleshooting with this device. This project was supported by MIRA grant (5R35GM137895) and an HMS Junior Faculty Armenise grant to MB. Marc W. Kirschner was supported by NIA R01AG073341 and MIRA R35GM145248. N.C.B was supported by following grants, National Science Foundation Graduate Research Fellowship Program (DGE 2140743) and Systems, Synthetic, and Quantitative Biology Training grant award (T32GM135014). Any opinions, findings, and conclusions or recommendations expressed in this material are those of the author(s) and do not necessarily reflect the views of the National Science Foundation.

