## supplementary_Information for "Bacterial cell wall biosynthesis is controlled by growth rate dependent modulation of turgor pressure in *E. coli*"

Mukherjee et al.

### Materials and methods

#### Bacterial strains used in this study

| Mentioned in manuscript as | Strain details | Comments |
|---|---|---|
| WT | NCM3722 | |
| MepS (YCE49) | NCM3722, pTet-TetR at ycad site, FRT-FRT:rrnBT:pTet:mepS, ΔmepH-FRT, ΔmepM::FRT | Tetracycline inducible mepS in mepH and mepM double knockout background |
| NQ393 | NCM3722 attB::Sp-lacIQ-tetR, ΔlacY, ΔgdhA, pLlac-O1-gltBD | IPTG inducible gltBD in gdhA knockout background. Used for titration of intracellular glutamate. Gift from Hwa lab |
| YCE44 | NCM3722, ΔfliC::FRT-FRT, glmS::PRNAI-mCherry1-11-mKate-T1 terminator-FRT Kan FRT::pstS | NCM 3722 strain with RNA1 promoter driven constitutive cytoplasmic RFP expression, used for turgor pressure measurement |
| NQ393_RFP | NQ393, glmS::PRNAI-mCherry1-11-mKate-T1 terminator-FRT Kan FRT::pstS | NQ393 strain with RNA1 promoter driving constitutive cytoplasmic RFP expression. Used for turgor pressure measurement |
| YCE15 | pZE1 ptetstab-LacZ in NCM3722 background, pTet-TetR at ycad site | Tetracycline inducible LacZ expression |
| YCE16 | pZE1pTetstab-(neg30) GFP in NCM3722, pTet-TetR at ycad site | Tetracycline inducible negatively supercharged GFP (-30) GFP (sub cloned from addgene plasmid #62936), in NCM3722 background, with pTet-TetR expressed from ycad site |
| YCE17 | pZE1 pTetstab-(pos36) GFP in NCM3722, NCM3722, pTet-TetR at ycad site | Tetracycline inducible positively supercharged GFP (+36) GFP (sub cloned from addgene plasmid #62937), in NCM3722 background, with pTet-TetR expressed from ycad site |

### Bacterial strain construction

**YCE49 strain construction:** NCM3722, pTet-TetR at ycad site, FRT-FRT:rrnBT:pTet:mepS, ∆mepH-FRT, ∆mepM::FRT

The NCM 3722 strain with the titratable pTet-TetR at ycad site was obtained as a gift from Hwa lab. This strain was used as the background strain for the following constructions:

For integrating the Tet promotor to the mepS gene chromosomally, the pkD13 plasmid was used for amplifying the pTet promotor with primers that also contain the mepS target region for replacing the mepS gene's own chromosomal promotor. The PCR product was sequenced before chromosomally integrated into the target strain using pKD46 recombineering method and the kan cassette was flipped out by the one-step inactivation method as described by Datsenko and Wanner, 2000 [1].

The mepH and mepM gene were then knocked out sequentially with the *mepHΔ* and *mepMΔ* p1 lysate generated from Kieo collection following methods described by Thomason L.C. et al, 2007 [2]; Baba T., et al, 2006 [3] . The kan cassette was then flipped out by the one-step inactivation method as described by Datsenko and Wanner, 2000 [1].

The generated strain YCE49 is kept in growth medium with CTC (50 ng/ml) to maintain the growth as the three genes (mepS, mepH and mepM) are together essential.

**NQ393_RFP strain construction:** RNA1 promoter driving mCherry-mKate::Kanamycin$^r$ was transferred to NQ393 strain by P1 transduction protocol modified from Thomason et. Al[2]. We obtained the donor strain (MG1655 glmS::PRNAI-mCherry1-11-mKate-T1 terminator-FRT Kan FRT::pstS) as a kind gift from Somenath Bakshi from Johan Paulsson lab.

**YCE44 strain construction:** Same as mentioned before RNA1 promoter driving mCherry-mKate :: Kanamycin$^r$ was transferred to NCM3722 strain with ΔfliC strain by P1 transduction. ΔfliC is less motile being a flagella knockout and helpful for microfluidics and microscopy experiment and does not cause any growth defect.

**YCE15, YCE16 and YCE17 strain construction:** These strains express different proteins from a tetracycline inducible plasmid (pZE1) [4] in the background of NCM3722 expressing tetracycline promoter driven Tet$^R$ from ycad site for linear induction of expression[5]. YCE15 expresses wildtype LacZ, YCE16 expresses negatively supercharged GFP (-30) and YCE17 expresses positively (+36) supercharged GFP[6,7]. The plasmid pZE1 expressing LacZ was previously published[4,8] and used as the starting point. Negatively and positively supercharged GFP plasmids were obtained from addgene (GFP (-30) #62936, GFP (+36) #62937) and cloned into the plasmid pZE1 ptetstab tetracycline inducible vector[4]. Plasmids were then transformed into an NCM3722 background expressing tetracycline promoter driving Tet$^R$ from ycad site [5].

### Bacterial culture methods

**Media formulation:** For most of the bacterial culture mentioned in this study, we used single Nitrogen and Carbon source containing (N+C+) minimal media following the protocol mentioned in previously published study [9]. Briefly, a 4X N-C- stock salt solution was prepared, following the recipe mentioned here:

1L Potassium free 4X N-C- salt buffer solution was prepared as followed:

| Species | Formula Weight | Concentration in 4X | g in 1l |
|---|---|---|---|
| $K_2SO_4$ | 142.04 | 23 | 3.26 |
| $K_2HPO_4$ | 141.96 | 310 | 44.01 |
| $KH_2PO_4$ | 119.98 | 138 | 16.57 |
| $MgSO_4$ | 120.37 | 1.6 | .192 |
| NaCl | 58.44 | 171.1 | 10 |

Using this 4X N-C- stock solution, bacterial growth media is prepared, in which 4XN-C- salt solution was diluted to achieve 1X concentration and the final media is supplemented with desired amount of carbon and 20 mM $NH_4Cl$. We kept the number of carbon atoms same in per unit volume of media, thus any 2-carbon source molecule (such as acetate) is added to obtain a final concentration of 60 mM, 3 carbon source molecules (Such as Glycerol) is added to obtain final concentration of 40 mM, and following the same principle 6 carbon atom containing sources (such as glucose) were added to obtain final concentration of 20 mM.

**Chemicals used in this study:** $K_2SO_4$ (VWR BDH Chemicals, Catalog #: BDH4618-500G), $K_2HPO_4$ (VWR BDH Chemicals, Catalog #: BDH9266-2.5KG), $KH_2PO_4$ (VWR BDH Chemicals, BDH9268-2.5KG), $MgSO_4$ (Sigma Aldrich, M7506-1KG), NaCl (Sigma Aldrich, S7653-1KG)

Carbon sources: D(+) Mannose (Sigma Aldrich, M6020-25G, Lot #BCBV4824), Glucose (Sigma, G5146), Sodium Acetate (Sigma S2889), Glycerol (VWR BDHH1172-1LP)

All media were filtered through 0.22µm PES vacuum filter. This media formulation was used for most of the experiments mentioned in this study, unless otherwise mentioned.

**General growth condition and growth rate determination:** For each growth condition, a single bacterial colony was inoculated in 5 ml LB broth culture and incubated in a shaker air incubator (Infros HT) set at 37˚C with 200 rpm orbital shaking. After few hours of growth, 1% of inoculum was transferred to glass test tubes containing 5 ml N+C+ minimal media (with desired carbon source) and incubated overnight in the same air incubator mentioned above.

To obtain exponentially growing cultures for experimentation, next morning we took out 1% inoculum from the overnight culture and inoculated in 5 ml fresh N+C+ minimal media in glass test tubes. Freshly inoculated tubes were incubated in a shaking water bath (benchmark scientific), set at 37˚ C with 200 rpm. For optical density measurement and growth curve determination, we took out samples from the growing cultures at intermittent time and measured $OD_{600}$ using a spectrophotometer (Thermo scientific, Genesys 30 visible spectrophotometer).

Experiments were performed in exponential range of bacterial growth (in between $OD_{600}$ 0.2 to 0.5), and growth rates were obtained by plotting the $OD_{600}$ as a function of elapsed time using semi-log scale.
For EZRDM media formulation mentioned in this study, we used the EZRDM media preparation kit from Teknova (Teknova: EZ Rich Defined Medium Kit, VWR, supplier no. M2105, VWR catalog number: 200059-658) and media was prepared following the manufacturer's guideline, and desired amount of carbon sources were added as mentioned.

To grow the NQ393 and MepS strains, we added required inducers (IPTG and CTC, chlortetracycline) to the initial LB culture. Inducers were also added to the overnight culture in minimal media and the freshly inoculated exponential growth cultures at desired induction level. This is very important to grow these two strains with respective inducers prior to beginning the exponential culture, as expression of these genes are required for growth. In absence of inducers, strains tend to mutate, and a fitter mutant version takes over the culture overnight, defying the purpose of the experiment. This is important to check the growth rates of these 2 strains in exponentially growing culture, to ensure the strains are not mutated and growing at a growth rate similar to wildtype. Also, both IPTG and CTC can give some variability in growth rates for the same inducer concentration on a day-to-day basis. This depends on the age of the stock solution and also on the manufacturer from which the chemical was obtained. Thus, we grouped inducer-dependent experiments primarily based on growth rates rather than inducer concentrations.

**Growth in chemostat:** For the chemostat setup, we used 'Chi.Bio' bioreactors [10], which are now commercially available to purchase from https://chi.bio/. These bioreactors use small volumes of culture (20 ml) and can keep culture temperature at constant 37˚ C with magnetic stirrer based continuous stirring. To run these bioreactors in chemostat mode, we capped the culture volume by placing a vacuum suction needle hanging at constant height within the bioreactor while fresh media was replenished at a constant rate in the culture by a peristaltic pump (Langer instruments). Chemostat cultures need to run out of the provided carbon source, and the growth and doubling time of the culture is determined by the rate at which new media is replenished. For chemostat experiments, all steps of standard growth were followed up to overnight culture in minimal media (N+C+Glucose), but next morning cells were inoculated in minimal media containing 0.01% glucose (final concentration). We determined previously that 0.01% glucose allows cells to grow up to 0.1 OD before going to diauxic conditions. First, we allowed cells to grow in flask and incubated in air incubator for some time, then 20 ml of the culture was transferred to Chi.Bio bio reactor vials. These bio reactors provided readings of culture turbidity, and when culture turbidity saturates, peristaltic pumps were turned on and bioreactors were allowed to run for long time to allow cells to adjust to slow growth rate conditions. Cell samples were taken from each bioreactor using long pipette tips and imaged by quantitative phase microscopy.

### Potassium measurement

Intracellular potassium ion measurements were done using ICP-MS method, following a protocol modified from the recent publication of Hwa group [11]. For this experiment, we have replaced all potassium ions in our N-C- salt buffer stock solution with sodium salts. 1L potassium-free 4X N-C- salt buffer solution was prepared as following:

| Species | Formula weight | Concentration in 4X (mM) | Grams added in 1L |
|---|---|---|---|
| $Na_2SO_4$ | 142.04 | 23 | 3.26 |
| $Na_2HPO_4$ | 141.96 | 310 | 44.01 |
| $NaH_2PO_4$ | 119.98 | 138 | 16.57 |
| $MgSO_4$ | 120.37 | 1.6 | 0.192 |
| NaCl | 58.44 | 171.1 | 10 |

From this 4X N-C- stock salt solution, we prepared the low potassium N+C+. minimal media, following the same protocol as before. 1X concentration and the final media was supplemented with desired amount of carbon as before, 20 mM $NH_4Cl$, and 0.1 mM (final concentration) potassium. Potassium was added from a 1M $KH_2PO_4$ stock solution.

Low potassium rich defined media was made modifying the recipe made by Neidhardt et al. [12] , where all potassium salts are replaced with sodium salts. To prepare this media, we prepared micronutrient stock and sodium salt supplemented 10X MOPS buffer in the lab, and used other components (where potassium replacement was not required) from the Teknova EZ rich defined media kit.

50ml micronutrient sock solution was prepared as following:

| Species | Chemical formula | Formula weight | Grams per 50 ml |
|---|---|---|---|
| Ammonium molybdate tetrahydrate | $(NH_4)_6Mo_7O_{24} \cdot 4H_2O$ | 1235.9 | 0.009 |
| Boric acid | $H_3BO_3$ | 61.83 | 0.062 |
| Cobalt chloride | $CoCl_2$ | 237.9 | 0.018 |
| Cupric sulfate | $CuSO_4$ | 249.7 | 0.006 |
| Manganese chloride | $MnCl_2$ | 197.9 | 0.040 |
| Zinc sulfate | $ZnSO_4$ | 287.5 | 0.007 |

10X MOPS buffer (potassium replaced with sodium) preparation:

First MOPS and Tricine mixed in 300 ml MilliQ water:

| Species | Formula weight | Grams |
|---|---|---|
| MOPS | 209.3 | 83.72 |
| Tricine | 179.2 | 7.17 |

MOPS buffer pH was adjusted to 7.4 with 10M NaOH and entire buffer volume was brought to 440 ml, to which 10 ml of freshly prepared 0.01M $FeSO_4$,$7H_2O$ was added. The following components were added to MOPS/Tricine/$FeSO_4$ solution in following order:

| Species | Volume added |
|---|---|
| 1.9 M $NH_4Cl$ | 50 ml |
| 0.276 M $Na_2SO_4$ | 10 ml |
| 0.02 M $CaCl_2$, $2H_2O$ | 0.25ml |
| 2.5 M $MgCl_2$ | 2.1 ml |
| 5 M NaCl | 100 ml |
| Micronutrient sock | 0.2 ml |
| MilliQ water | 387 ml |

Now 1L low potassium rich defined media was used in following way:

| Component | Volume | Source |
|---|---|---|
| 10X MOPS buffer | 100 ml | Prepared in lab |
| 0.132 M $Na_2HPO_4$ | 10 ml | |
| 1M $KH_2PO_4$ | 100µl | |
| 1M Glycerol | 40 ml | |
| 10X ACGU solution | 100 ml | Teknova kit |
| 5X EZ supplement | 200 ml | |

Potassium was added to achieve final concentration of 0.1 mM and 40 mM glycerol (final concentration) was maintained in this low potassium rich defined media. In low potassium conditions, glucose was not used as a carbon source as glucose transport has potassium dependency and growth rate slows.

For this experiment all glassware and plasticware were pre-rinsed with 1% (V/V) sub-boiling-$HNO_3$ (prepared from High Purity Standards sub-boiling $HNO_3$, Cat. SB-HNO3-250, Lot. 2218110-250), followed by washing with excess MilliQ water.

2 colonies of NCM3722 WT strain were inoculated in LB for few hours and from LB culture 1% inoculum was transferred to low potassium minimal media (with 20 mM mannose, 20 mM mannitol, 40 mM glycerol,

separately) or low potassium rich defined media with 40 mM glycerol. Freshly inoculated tubes were incubated overnight in shaker incubator.

Next morning, 1% inoculum from different overnight cultures were transferred to 100 ml fresh low potassium media in 500 ml baffled conical flask, which were incubated in the shaking air incubator. 200 µl samples were drawn out intermittently from the shaking cultures to measure $OD_{600.}$ For potassium measurements, 10 ml sample drawn out from the exponentially growing culture. Exact optical density of the culture and volume of the sample taken were noted down to aid normalization at the time of intracellular potassium concentration calculation.

After drawing the samples from exponentially growing cultures, cells were rapidly trapped on a filter paper (using a 0.22 µm filter placed in a glass vacuum filtration device). Cells were washed with osmotically balanced water (7% PEG 200 V/V in water). This washing step is necessary to remove extracellular potassium trapped on the filter paper. We ensured that the exact same volume of osmotically balanced water is used for each sample to minimize the error between samples. The protocol mentioned by Hwa group used low potassium media (carbon free) as a washing agent. As the low potassium media still contains potassium, we decided to use osmotically balanced water for washing. The washing step causes loss of some intracellular ions which cannot be avoided, but keeping the wash volume and time of wash same for all samples help in minimizing this systematic error.

After washing, filter paper was taken out and rapidly submerged in 10 ml of 1% sub-boiling nitric acid (High Purity Standards) kept in a 50 ml falcon, and kept for several hours in cold room to ensure complete lysis of bacterial cells in nitric acid. For intracellular potassium measurement, 1 ml of each sample were taken and further diluted in 9 ml 1% nitric acid.

The samples were analyzed in a solution-nebulization ICP-MS (SN-ICP-MS) at Harvard University Solid Earth Geochemistry lab using a Thermo Scientific X series ICP-MS set up.

We prepared a standard curve by running a standard of known potassium concentrations in the ICP-MS and potassium concentration in the samples were calculated by extrapolation from the standard curve. For standard curve preparation, we obtained dissolved metal standards (10 ppm each metal, 40 metal ions, dissolved in 1% nitric acid, High Purity Standards) and standards of known concentration were prepared from the stock solution by dissolving the 10-ppm standard solution in 1% nitric acid. To check consistency of measurements between samples, known amount of Rhodium (High Purity Standards) were added in all samples and standards.

To obtain potassium concentration per OD•ml , we first calculated potassium concentration in PPM value in each sample by extrapolation from standard curve. Each sample that was run in ICP MS was diluted 10-fold, so each PPM value was multiplied by 10, then PPM value is converted to mM concentration by multiplying with 0.02564 (PPM to mM conversion factor for K).

To obtain net potassium amount of each sample from potassium concentration, we multiplied the potassium concentration with total lysate volume (10 ml), and then by 0.001 to obtain mmol amount of K in each sample. Now, mMol amount of potassium in each sample was plotted as a function of [OD•ml] of sample taken (see Supplementary Fig. 10), and slope of the linear fit represents mmol/[OD•ml] of potassium for each growth rate condition.

### Turgor pressure measurement

To measure intracellular turgor pressure, exponentially growing bacteria on sudden osmotic shock experience detachment of plasma membrane at its poles from the cell wall and enter a state of plasmolysis. For bacterial cells that enter plasmolysis, the shrinkage of the volume is proportional to external osmotic pressure applied. For turgor pressure measurement, NCM3722 cells constitutively expressing cytoplasmic RFP were grown in a shaking water bath in N+C+ minimal media containing different carbon sources. Exponentially growing cells were loaded in Millipore Cell-Asic Onyx B01 (compatible for bacteria plates) following the manufacturer's protocol. Fresh minimal media (with the same carbon source in which cells were growing) were loaded in one of the media flow chambers of the microfluidics plate. In other flow chambers, we loaded osmotic shock media with different amounts of external osmolarity. To prepare osmotic shock media, we used 4X N-C- slats as a base and first added the same carbon source in which cells are exponentially grown, then sucrose was added to increase the osmolarity to the desired level. In this way, we kept all the salt and carbon concentration in osmotic shock media the same as in growth media, and the amount of added sucrose accounts for the enhanced external osmolarity. After loading cells and media on the microfluidics plate, we mounted the plate on inverted epi-fluorescence microscope stage (Nikon TiE) and started flowing the growth media to ensure cells are in exponential growth phase. We used a microscope with fitted environmental chamber and that allowed us to keep the plate temperature at 37˚ C. For proper turgor pressure measurement, it is extremely important to keep the cells in growth phase, as only exponentially growing cells exhibit turgor pressure. We imaged the cells using a 100X oil objective lens with phase contrast compatibility and time-lapse images were recorded in phase contrast and epi-fluorescence RFP channel. We flowed growth media for certain time, then rapid osmic shock was induced by flowing osmotic shock media from the chamber in which it was loaded previously, following the manufacturer's protocol. On osmotic shock, cells entered in plasmolysis, and cell images were recorded in the time-lapse movie.

To calculate turgor pressure, length of cell cytoplasm immediately before the shock ($L_0$) and the minimum length of cytoplasm after shock ($L_1$) was measured, and ratio of $L_1/L_0$ was calculated for each osmotic shock condition. Length of the cytoplasm is proportional to the volume of the cell cytoplasm and a measurable parameter to represent volume. Averaged ratio of lengths was then plotted as a function of inverse of external osmolarity. To obtain turgor pressure in each growth condition, we fit the data points to the Boyle-van't Hoff equation[13], and the y intercept of the fit line represents the incompressible volume offset.

### Quantitative Phase Microscopy

Quantitative phase imaging was performed using a commercial quadriwave lateral shearing interferometry (QWLSI [14]) wavefront sensor (Phasics, SID4BIO) attached to an inverted brightfield microscope (Nikon Eclipse Ti). All imaging was performed at 300x magnification using an 100x oil immersion objective lens (Nikon CFI Plan Apo Lambda) and additional 2x magnifier inserted after the tube lens. For the additional 3x magnification, the 1.5x auxiliary magnifier of Nikon microscope was engaged and an additional 2x magnifying lens was attached in front of the wavefront sensor. A halogen lamp was used for trans-illumination. Blue color filter was inserted in the illumination path to maximize the optical resolution. The condenser lens was positioned for the Koehler illumination. Aperture diaphragm was closed to the minimum position to create a plane wave at the sample plane.

To ensure bacterial cells are imaged in best possible physiological condition, we always set up the shaking water bath containing culture tubes, and spectrophotometer for OD measurement next to the microscope room. To image cells in the QPM microscope, we first created an imaging chamber by attaching Grace Bio-Labs secure seal precut imaging chambers with 120 µm height. These imaging chambers have double sided adhesive, and that allows attaching a 1.5 coverglass on top of the imaging chamber. 8µl bacterial culture was pipetted out from exponentially growing tubes and placed at the center of the imaging chamber, and a coverglass is placed on top to seal it. Then the slide was centrifuged in a plate centrifuge (Benchmark scientific) for 30-45 seconds ensuring enrichment of bacterial cells on the coverglass.

Immediately after spinning. The slide was placed in the motorized sample stage for image acquisition. The light transmitted through the sample was imaged on to QWLSI wavefront sensor (Phasics, SID4BIO). Focus position was determined by the minimum of the contrast of the brightfield mode image. The microscope was equipped with autofocus (Nikon, Perfect Focus) to maintain a constant focus during image acquisition. QWSLI interferograms were converted to quantitative phase images following the method reported in the prior publication[15]. Briefly, we employed this previously developed technique, computationally enhanced Quantitative Phase Microscopy (ceQPM), to measure the dry mass of individual cells. This method significantly improves the precision of dry mass quantification by incorporating two background subtraction steps. In the first step, the method calculates the median of the Fourier harmonics from multiple fields of view to synthesize a common reference wavefront, which is then subtracted from the sample wavefront. This subtraction yields the net phase shift resulting from the refractive index difference between the cell and the surrounding medium. In the second step, the method utilizes three-dimensional "thin plate" interpolation to effectively fit the residual wavefront fluctuations caused by variations in cover glass thickness and vibrations, thereby further leveling the background. By applying this two-step background subtraction method, we have successfully reduced the quantification error in dry mass measurement to below the 2% margin.

### Image processing and dry mass density calculation

In a quantitative phase image, pixel values are proportional to z-projection of biomass. (Fig. S2A) After background removal following the steps mentioned above, single cell images were cropped out using thresholding. At this step, we set a low threshold and also dialated the cropping mask to ensure entire cells are taken and there is no loss of biomass from single cell images. Cropped out single cell images were centered on a zero padded background.

QPM images are extremely accurate representation of the biomass, but to obtain density, bacterial volume must be calculated accurately. To do so, we simulated QPM image for each bacterium, assuming a bacterium is composed of a cylinder and two hemispheres at the end of the cylinder (Fig. S3). In this way, a bacterial QPM image can be simulated to obtain the precise length and width parameter values, which will ensure accurate volume calculation. For each real bacterial QPM image, we matched it to a simulated QPM image, and calculated the mean squared error (MSE) between the two images. To find best match for each real QPM image, we iteratively generated multiple simulated images and calculated mean squared error between the real and the simulated image at each iteration. We stopped the iterative simulation process when the minima of mean squared error was achieved for a particular real bacterial image, and volume of the cell was calculated from the length and width parameters of the best matched simulated cell. Bacterial biomass is always calculated from the real QPM image. Details of the bacterial cell simulation and density calculation by iterative matching with simulation is described in the following sections.

### Simulating QPM image of bacterial cell

**Computation of the optical field scattered by a refractive object:** Our goal is to calculate the scattering of a plane monochromatic field $u_i(\vec{r}) = \exp(ik_0 z)$ by a model *E. coli* described in Fig. S3. The resulting field $u(\vec{r}) = u_i(\vec{r}) + u_s(\vec{r})$, where $u_s(\vec{r})$ is the scattered field, satisfies the Helmholtz equation as per Wolf 1969 [16].

$$\nabla^2 u(\vec{r}) + {k_0}^2 n^2(\vec{r}) u(\vec{r}) = 0, \quad \text{[M1]}$$

where $k_0 = \frac{2\pi n_0}{\lambda}$ and $n(\vec{r})$ describes the relative refractive index to $n_0$. Since the plane wave incident field satisfies $\nabla^2 u_i(\vec{r}) + {k_0}^2 u_i(\vec{r}) = 0$, Eq. [M1] can be written as

$$\nabla^2 u_s(\vec{r}) + {k_0}^2 u_s(\vec{r}) = f(\vec{r}) u(\vec{r}), \qquad \text{[M2]}$$

where the scattering potential $f(\vec{r}) = {k_0}^2\left(1 - n^2(\vec{r})\right)$ is zero outside of the cell. Under the first-Born approximation, the solution to Eq. [M2] is given by:

$$u_s(\vec{r}) = -\frac{1}{4\pi} \iiint f(\vec{r}') u_i(\vec{r}') G(|\vec{r} - \vec{r}'|) d\vec{r}' \ , \qquad \text{[M3]}$$

where $G(r) = \exp(ik_0 r)/r$. We substitute the Green's function with the following representation:

$$G(x,y,z) = \iint \frac{i}{\lambda^2\sqrt{1/\lambda^2 - U^2 - V^2}} e^{i2\pi\left(Ux+Vy+(\sqrt{1/\lambda^2-U^2-V^2})z\right)} dUdV, \qquad \text{[M4]}$$

and the incident field $u_i(\vec{r}) = \exp(ik_0 z)$ in Eq. [M3] to obtain:

$$u_s(\vec{r}) = \frac{1}{i4\pi} \iint \frac{\tilde{f}(U,V,\sqrt{1/\lambda^2-U^2-V^2}-1/\lambda)}{\lambda^2\sqrt{1/\lambda^2-U^2-V^2}} e^{i2\pi\left(Ux+Vy+(\sqrt{1/\lambda^2-U^2-V^2})z\right)} dUdV. \qquad \text{[M5]}$$

in which $\tilde{f}$ is the Fourier transform of $f$. We rewrite Eq.[S19] using $W = \sqrt{1/\lambda^2 - U^2 - V^2} - 1/\lambda$:

$$u_s(\vec{r}) = \frac{1}{i4\pi} \iint \frac{\tilde{f}(U,V,W)}{\lambda^2(W+1/\lambda)} e^{i2\pi(Ux+Vy+(W+1/\lambda)z)} dUdV \ . \qquad \text{[M6]}$$

At z=0, we take the inverse Fourier transform with respect to x and y:

$$\tilde{u}_s(U,V) = \frac{\tilde{f}(U,V,W)}{i4\pi(W+1/\lambda)}. \qquad \text{[M7]}$$

Eq. [M7] allows us to calculate the scattered field from the scattering potential of the *E. coli* model as per Sung et al. 2013 [17].

**Computation of the scattering potential of an object with a uniform refractive index:** Given the scattering potential $f(\vec{r}) = \left(\frac{2\pi n_0}{\lambda}\right)^2 (1 - n(\vec{r})^2)$ of a cell with a uniform relative refractive index $n(\vec{r}) = n_{cell}/n_0$ for $\vec{r}$ inside the cell and $n(\vec{r})$=1 for $\vec{r}$ ouside the cell, the Fourier transform of the scattering potential is

$$\begin{aligned}\tilde{f}(U,V,W) &= \iiint f(x,y,z) e^{-i2\pi(Ux+Vy+Wz)} dxdydz \\ &= Q_s \iiint_{V_{cell}} e^{-i2\pi(Ux+Vy+Wz)} dxdydz,\end{aligned} \qquad \text{[M8]}$$

where $Q_s = \left(\frac{2\pi n_0}{\lambda}\right)^2 \left(1 - \left(\frac{n_{cell}}{n_0}\right)^2\right)$.

Following Sung et al 2015 [18], we define the following vector field $\vec{F}$ as

$$\vec{F}(x,y,z;k_x,k_y,k_z) = \frac{1}{3} e^{-i2\pi(k_x x+k_y y+k_z z)} \begin{bmatrix} x sinc(k_x x) e^{i\pi k_x x}\hat{\imath} \\ +y sinc(k_y y) e^{i\pi k_y y}\hat{\jmath} \\ +z sinc(k_z z) e^{i\pi k_z z}\hat{k} \end{bmatrix}, \qquad \text{[M9]}$$

where $sinc(t) = sin(\pi t)/(\pi t)$.

$\vec{F}$ is continuously-differentiable and satisfies the condition

$$\nabla \cdot \vec{F} = e^{-i2\pi(k_x x+k_y y+k_z z)}. \qquad \text{[M10]}$$

Combining these equations, Eq. [M9] can be expressed as a surface integral by the divergence theorem

$$\tilde{f}(U,V,W) = Q_s \iiint_{V_{cell}} \nabla \cdot \vec{F} dxdydz = Q_s \oiint_{S_{cell}} \vec{F} \cdot d\vec{S} \,. \qquad \text{[M11]}$$

Which can be computed using the non-uniform rational B-splines (NURBS) coordinates $u$ and $v$ as per Sung et al 2015 [18] :

$$\oiint_{S_{cell}} \vec{F}(x,y,z) \cdot d\vec{S} = \oiint_{S_{cell}} \vec{F}(u,v) \cdot \left(\partial\vec{S}/\partial u \times \partial\vec{S}/\partial v\right) dudv. \qquad \text{[M12]}$$

The surface normal vectors and coordinates were obtained following the algorithm described in Piegl and Tiller [19] using the code by D. M. Spink (https://www.mathworks.com/matlabcentral/fileexchange/26390-nurbs-toolbox-by-d-m-spink) in MATLAB (MathWorks, Inc.).

Eq. [M11] and Eq. [M12] allow us to obtain scattered field $u_s(\vec{r})$ from Eq. [M7]. The phase of the field in real space is the desired quantitative phase image of the cell.

**Dry mass density calculation by iterative simulation match:** We performed iterative matching of the real QPM image with simulated QPM images generated by the method described above using a custom MATLAB script. First, a single real bacterial cell image was cropped out from QPM data and centered horizontally on a zero-padded background. The real image is computationally up sampled 4 times to reduce pixel size, which enhances the accuracy of the simulation. By thresholding and fitting a bounding box, we roughly measured the length and width of the bacteria (estimated from real microscopic image by a custom-built Fiji script as a preprocessing step), and this information is used as initial parameters for the simulation. In the simulation, we iterated four parameters (length, width, refractive index, and rotation ($R_y$) of the simulated cell). (Fig. S3A-B) We used the MATLAB Optimization Toolbox function *fminsearch*, a nonlinear programming solver to find the minimum of our objective function. For each iteration, mean squared error (MSE) is calculated between the real image and the simulated image, and the simulation is ended when a MSE minima is reached. Cell volume is calculated from the length and width parameters of the matched simulated cell. (Fig. S3A) In QPM microscopy, the areal integral of quantitative phase over the cell area is proportional to cellular dry mass, and the refractive index increment value serves as the proportionality constant. (Fig. S2B) We used a refractive index increment value of 0.18 ml/g which is a commonly used value for average protoplasm [20,21].

### Threshold Iterative Volume (TIV) Determination

We developed a second method for determining bacterial volumes that has advantages in certain conditions because it only assumes axial symmetry of bacteria. QPM measurements provide us with a 2-dimensional projection of the dry mass distribution of the cell. We focus on cells that are lying flat on the coverslip. By setting a low threshold, we determine which pixels contribute to the total mass of the cell. The total mass of the cell can then be calculated by summing up the individual mass contributions from the individual pixels within the cell contour. By exploiting the axial symmetry of rod-shaped bacteria like *E. coli*, we can calculate the volume of the bacterium by rotation of the cellular contour around the symmetry axis. The symmetry axis is determined by fitting the smallest possible rectangular box that fully contains the cell contour.

The major difficulty with this approach for volume determination is selecting the correct threshold for determining the cell contour determining the cell volume. Due to the cylindrical shape of the cell and diffraction limit of light, there is no straightforward way to know the true boundary of the cell. However, small changes in this contour will have a rather large effects on the volume of the cell and the calculated dry mass density. To resolve this problem, we make use of the axial symmetry of rod-shaped bacteria: The width of the cell, determined from the contour, is equal to the length of the light path for groups of pixels along the symmetry axis in the center of the cell. Therefore, the phase shift of the light for these pixels is proportional to the width of the cell multiplied by the dry mass density of the cell. Assuming a homogenous mass distribution, this dry mass density should be equal to the dry mass density calculated from the total mass of the cell, divided by the cell volume determined from the contour using the correct cell volume threshold. Hence, when the volume threshold for determining the cell volume contour is correct, these measurements should be self-consistent. Therefore, in our method, we iteratively increase the volume threshold for the cell volume contour, until the two determinations of dry mass density are self-consistent.

While there is a small shift in absolute values of dry mass densities from the two methods, the threshold iterative volume (TIV) method shows good agreement with the QPM simulation methods and recapitulates changes that we found using the simulation method. The advantage of the threshold iterative volume (TIV) method as compared to the QPM simulation method is that only axial symmetry is required for the TIV method to work. Thus, unlike the QPM simulation, the shape of the bacterium does not need to be close to a perfect capsule shape. For very long cells, such as those with useless protein overexpression, we observe changes in cell width along the length of the bacterium, making the QPM simulation method unsuitable, whereas the self-consistent TIV method still works.

### Supplementary Note 1: Viscoelastic model of cell wall expansion

#### Model summary

In one dimension, the constitutive equation of a viscoelastic Maxwell fluid is given by

$$\eta \, \partial_x v \; = (1 + \tau \partial_t)\sigma, \qquad \text{[S1]}$$

where $v$ is the velocity of cell wall material, $\sigma$ is mechanical stress, $\eta$ is the effective bulk viscosity, and $\tau$ is the viscoelastic relaxation time[22]. On timescales shorter than $\tau$, the cell wall behaves as an elastic material with elastic modulus $E = \eta/\tau$. The timescale $\tau$ is determined by the rate of cell wall remodeling by endopeptidases $k_{hyd} \sim 1/\tau$. On longer timescales, the time derivative can be neglected, and the material behaves like a viscous fluid.

We approximate the geometry of the bacterium as a cylinder of length $L$ with the cylindrical axis along the $x$-axis. Cell width is twice the cell radius, denoted by $2r$. Turgor pressure $P_{turg}$ in the cytoplasm acts on the cross section of the cylinder, generating a force given by $r^2 \pi P_{turg}$, which is balanced by the tension in the cell wall along the $x$-axis. Assuming the tension is uniformly distributed around the cell circumference of length $2\pi r$, the cell wall tension is given by

$$\sigma = \frac{r^2 \pi P_{turg}}{2\pi r d} = \frac{r P_{turg}}{2d}, \qquad \text{[S2]}$$

where $d$ a dimensional constant reflecting the cell wall thickness. This cell wall tension drives the elongation of the bacterium and the elongation speed $\dot{L}$ can be determined by inserting Eq. [S2] into Eq. [S1] and integrating over the cell length along $x$ from 0 to $L$. Hence, the volume growth rate $\lambda_V$ of the bacterium is given by

$$\lambda_V = \frac{\dot{V}}{V} = \frac{\dot{L}}{L} = \frac{r P_{turg}}{2\eta d}. \qquad \text{[S3]}$$

#### Turgor pressure proportional to growth rate maintains constant biomass density across growth rates

During steady-state growth, the biomass density $\rho$ is constant and therefore the volume growth rate $\lambda_V = \dot{V}/V$ is equal to the biomass growth rate $\lambda_M = \dot{M}/M$:

$$\lambda_V = \lambda_M. \qquad \text{[S4]}$$

Assuming constant cell shape and cell wall properties, to satisfy Eq. [S4] and Eq. [S3] across different growth rates, turgor pressure must scale directly proportional to biomass growth rate:

$$P_{turg} \propto \lambda_M. \qquad \text{[S5]}$$

### Dry mass density decreases with cell width

An increase in cell width affects volume growth rate of the cell according to Eq. [S3] because turgor pressure acts on a larger cross section, resulting in higher expansive tension in the cell wall (see Eq. [S2]). If biomass growth rate and cell wall properties remain unchanged, Eq. [S3] dictates that a drop in turgor pressure must compensate for this larger cross section:

$$rP_{turg} = const. \qquad \text{[S6]}$$

Moreover, we assume that turgor pressure is proportional to biomass density according to the mechano-electro-osmotic model Eq. [4], main text:

$$P_{turg} \propto \rho. \qquad \text{[S7]}$$

Combining Eqs. [S6] and [S7], we obtain that dry mass density should be inversely proportional to cell width for perturbations of cell width that do not affect growth rate:

$$\rho \propto {}^{1}/_{r}. \qquad \text{[S8]}$$

### Dry mass density decreases with cell wall elasticity

The effective viscosity of the cell wall is given by $\eta = E\,\tau$. Therefore, decreasing the elastic modulus of the cell wall $E$ by treatment with beta-lactam antibiotics that block insertion of cell wall precursors and crosslinking of the cell wall network results in a lower viscosity and a faster volume expansion rate for a given turgor pressure. However, if growth rate is constant, the expression given by Eq. [S3] must be constant. Combining Eq. [S3] with Eqs. [S4] and [S7] and inserting the expression for the viscosity above, we obtain

$$\frac{r\,\rho}{2E\,\tau\,d} = const. \qquad \text{[S9]}$$

If the width of the cell remains unchanged, as we observe experimentally, the lower viscosity must be compensated by a drop in dry mass density, and we obtain

$$\rho \propto E. \qquad \text{[S10]}$$

### Downregulation of cell wall hydrolases increases cell width and dry mass density

The effective viscosity of the cell wall $\eta$ is determined by the product of the elastic modulus of the cell wall $E$ and the viscoelastic relaxation time $\tau$, which is determined by the rate of remodeling of the cell wall by hydrolases, denoted by $k_{hyd}$:

$$\tau \propto 1/k_{hyd}. \qquad \text{[S11]}$$

Hence, for a constant elastic modulus of the cell wall with saturating insertion of peptidoglycan, the viscosity is inversely related to the rate of endopeptidase activity

$$\eta \propto {}^{1}/_{k_{hyd}}. \qquad \text{[S12]}$$

Therefore, higher cell wall viscosity due to downregulation of endopeptidases directly affects volume growth rate according to Eq. [S3]. Combining Eqs. [S3], [S4], [S7] and [S12], we obtain a quantitative prediction for changing endopeptidase expression level:

$$\frac{\lambda}{r\,\rho} \propto k_{hyd}. \qquad \text{[S13]}$$

Note that in this situation growth rate changes due to molecular crowding as opposed to different substrates. Ribosome content is roughly constant and does not scale with growth rate in this condition, but only with biomass density.

#### Dependence of turgor on both biomass density and growth rate

Eq. [S5] and Eq. [S7] may appear at first glance to be in contradiction: how can turgor pressure be both proportional to growth rate and biomass density? However, this is precisely the dependence expected from the hypothesis introduced in Fig. 5 of the main text. If turgor pressure is determined by the charge concentration of intracellular RNA, mostly originating from ribosomal RNA, the turgor pressure is both proportional to growth rate in steady-state conditions because ribosomal concentrations increase with growth rate due to growth laws and proportional to biomass density because ribosomes are a component of biomass. Therefore, Eq. [S5] and Eq. [S7] can be combined:

$$P_{turg} \propto \rho\lambda_M. \qquad \text{[S14]}$$

### Supplementary Note 2: Electro-osmotic model of turgor pressure

This model is a straight-forward adaptation for bacteria, of a previously developed electro-osmotic model for mammalian cells[23]. This model combines flux balance of each ion species across the plasma membrane from diffusion, electrophoretic mobility, and active transport with charge neutrality of the cytoplasm and mechanical force balance of osmotic forces.

There are small modifications of the model for bacteria. Specifically, since there is no sodium-potassium-ATPase in *E. coli*, we assume individual pumps for every ion species. For simplicity, we assume a single Michaelis Menten function for each ion species. Based on the observed high abundance of intracellular potassium and low concentrations of all other ions compared to medium concentrations[24,26,28], we assume that potassium is actively imported, whereas all other ions are actively exported. Composition of the reference medium was based directly on our experimental growth medium for bacteria. Concentrations are given in the table below.

| **Ion** | **Concentration [mM]** |
|---|---|
| $Cl^-$ | 62.8 |
| $K^+$ | 201.7 |
| $Na^+$ | 42.8 |
| $SO_4^{2-}$ | 6.1 |
| $HPO_4^{2-}$ | 77.5 |
| $H_2PO_4^-$ | 34.5 |
| $NH_4^+$ | 20.0 |

### Supplementary Figures:

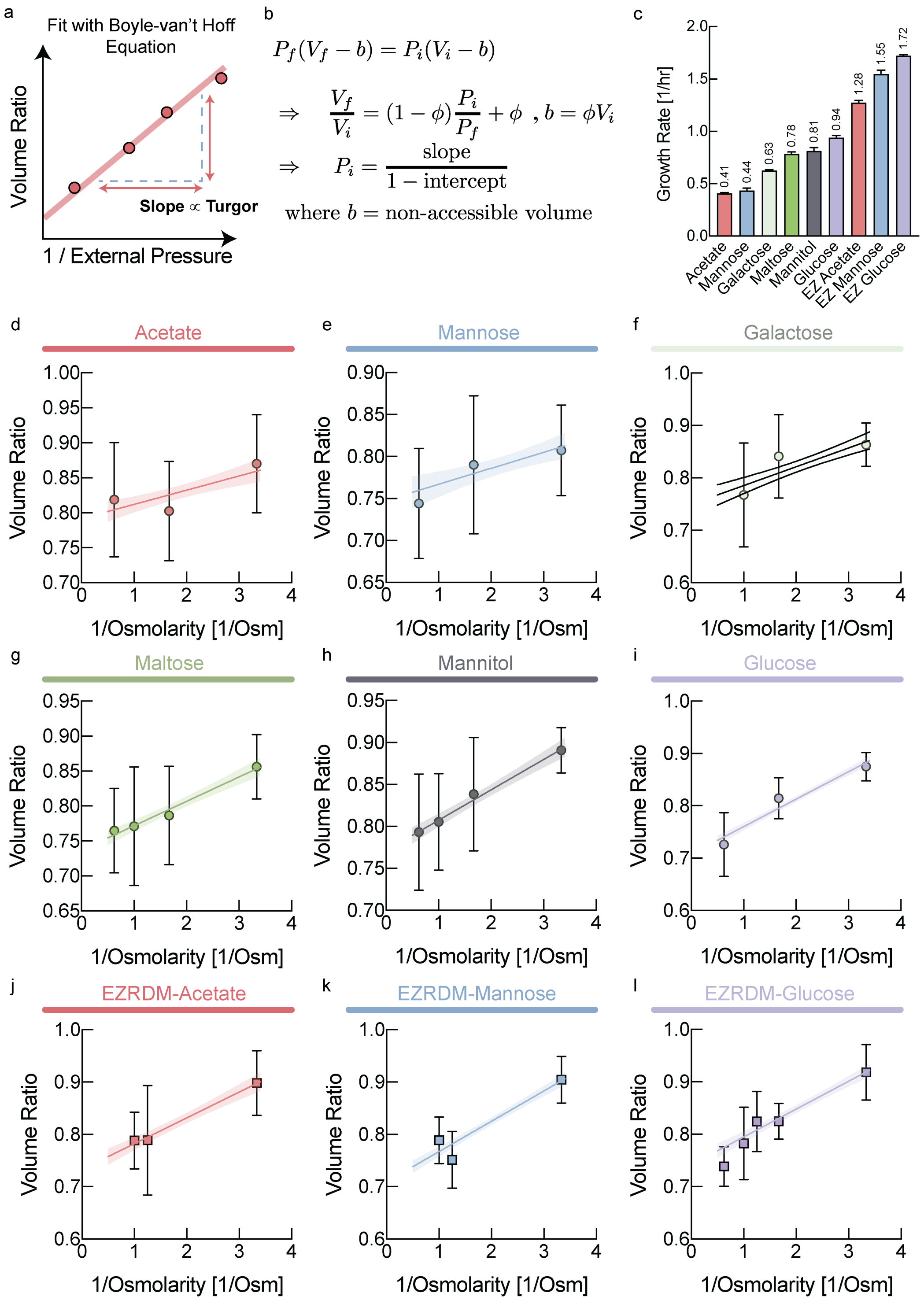

*__Fig. S1: Inferring turgor by inducing plasmolysis via osmotic shocks. a,__ Schematic diagram representing the method used to calculate the turgor pressure__.__ Ratio of cytoplasmic volume before*

*and after osmotic shocks plotted against inverse osmolarity of the applied osmotic shock.* ***b,*** *Equations used to determine turgor pressure. Slope of the curve, as described in panel A, fitted to Boyle-van't Hoff equation as described here.* ***c,*** *Growth rate of E. coli NCM3722 growing in different carbon sources, measured from the same samples that are used for turgor pressure measurement.* ***d-i,*** *Ratio of the cytoplasmic length (proportional to volume as we observe a roughly constant cell width). Individual data points represent biological replicates. Each data point results from averaging 13 randomly selected cells for this osmotic shock and error bars represent the resulting standard deviation. According to the Boyle-van't Hoff equation (see panel B), the slope of these curves is proportional to initial turgor pressure. The data in Fig. 1C of the main text was determined by fitting the slope as described above. The y-intercept reflects a phenomenological excluded volume (e.g. volume of biomass components).*

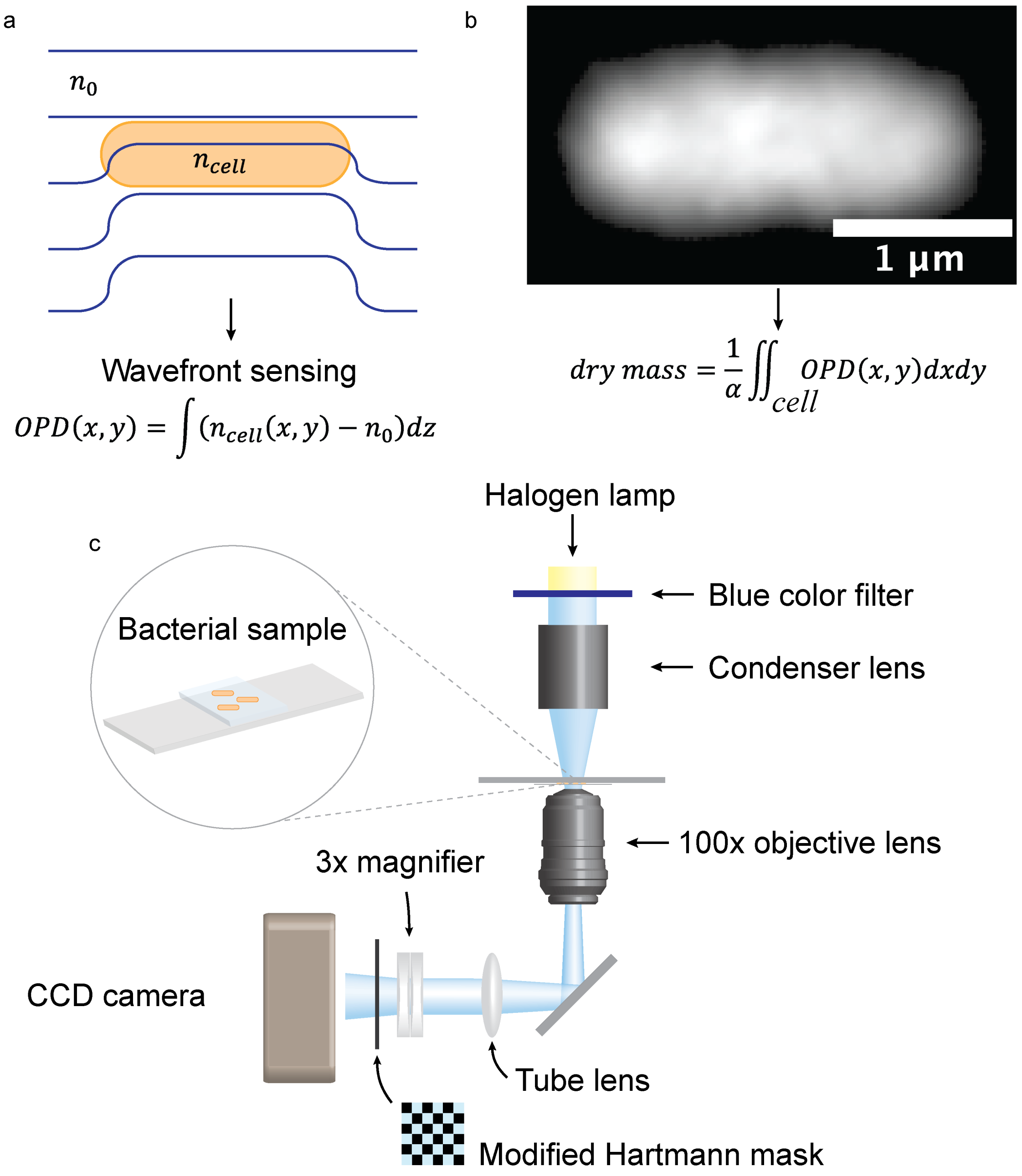

***Fig. S2: Quantitative phase microscopy (QPM). a**, E. coli cells were laid flat on a coverslip (not shown) and illuminated with a plane wave. The delay of the wavefront that traversed each cell was measured by a wavefront sensor to determine the optical path difference (OPD). OPD is determined by the refractive index of cell ($n_{cell}$) and media ($n_0$) and cell height. **b**, Dry mass of single e coli cell is proportional to the OPD integrated over the cell area. The average refractive index increment of cytoplasm (0.18 ml/g) was used for the proportionality constant ($\alpha$). **c,** Schematics of quantitative phase imaging microscope. Blue color filter was applied to halogen lamp light source to maximize the optical resolution. The condenser lens was put in the Koehler illumination position and aperture diaphragm (not shown) was closed to create a plane wave at the sample plane. Bacterial sample was placed on a slide glass and imaged through a coverslip by 100x oil immersion objective lens. The transmitted light was imaged on to QWLSI wavefront sensor (Phasics, SID4BIO) to acquire the quantitative phase image of the sample*

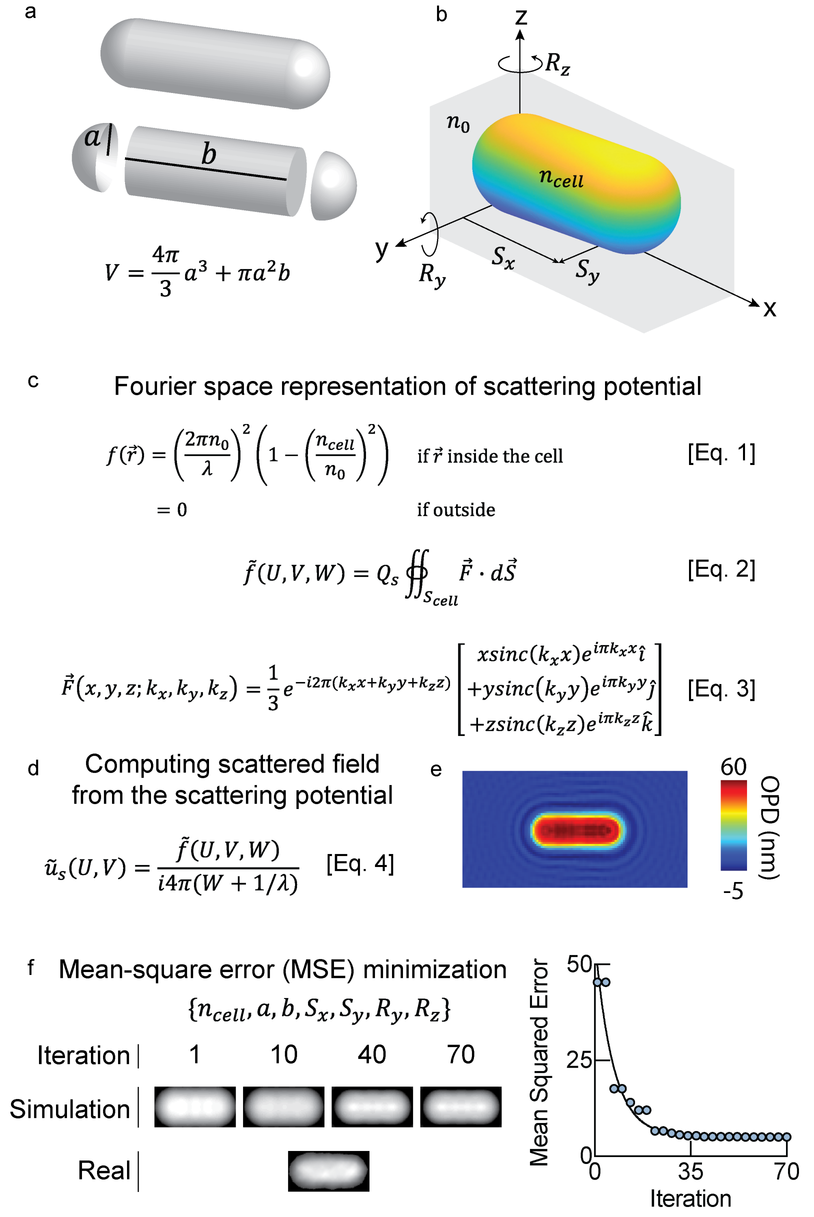

***Fig. S3: Simulation of QPM image for determining cell volume. a,*** *The optical model of an e coli for quantitative phase imaging simulation. A cell is in the shape of a cylinder with hemisphere caps at both ends and has a uniform refractive index ($n_{cell}$) inside.* ***b,*** *By centering and rotating the image to align the cell horizontally, two positional ($S_x$, $S_y$) and one rotational ($R_z$) degrees of freedom are removed. Z-translation is also removed as out-of-focus images are rejected. As a result, a single e coli cell in the imaging plane has a single rotational ($R_y$) degree of freedom.* ***c,*** *[Eq.1] Scattering potential $f$ of a cell with a uniform refractive index ($n_{cell}$) immersed in media ($n_0$) illuminated by light of wavelength $\lambda$. [Eq. 2] Fourier transform of the scattering potential can be expressed as a surface integral of a vector field $\vec{F}$ as defined in Sung et al. [Eq. 3] using the divergence theorem. $Q_s = \left(\frac{2\pi n_0}{\lambda}\right)^2 \left(1 - \left(\frac{n_{cell}}{n_0}\right)^2\right)$* ***d,*** *The light field scattered by a cell is related to the cell's scattering potential in Fourier domain as shown in Eq. 4. U and V are the Fourier conjugates of x and y, and $W = \sqrt{1/\lambda^2 - U^2 - V^2} - 1/\lambda$.* ***E,*** *The phase of the scattered field in image space provides the quantitative phase image in OPD.*

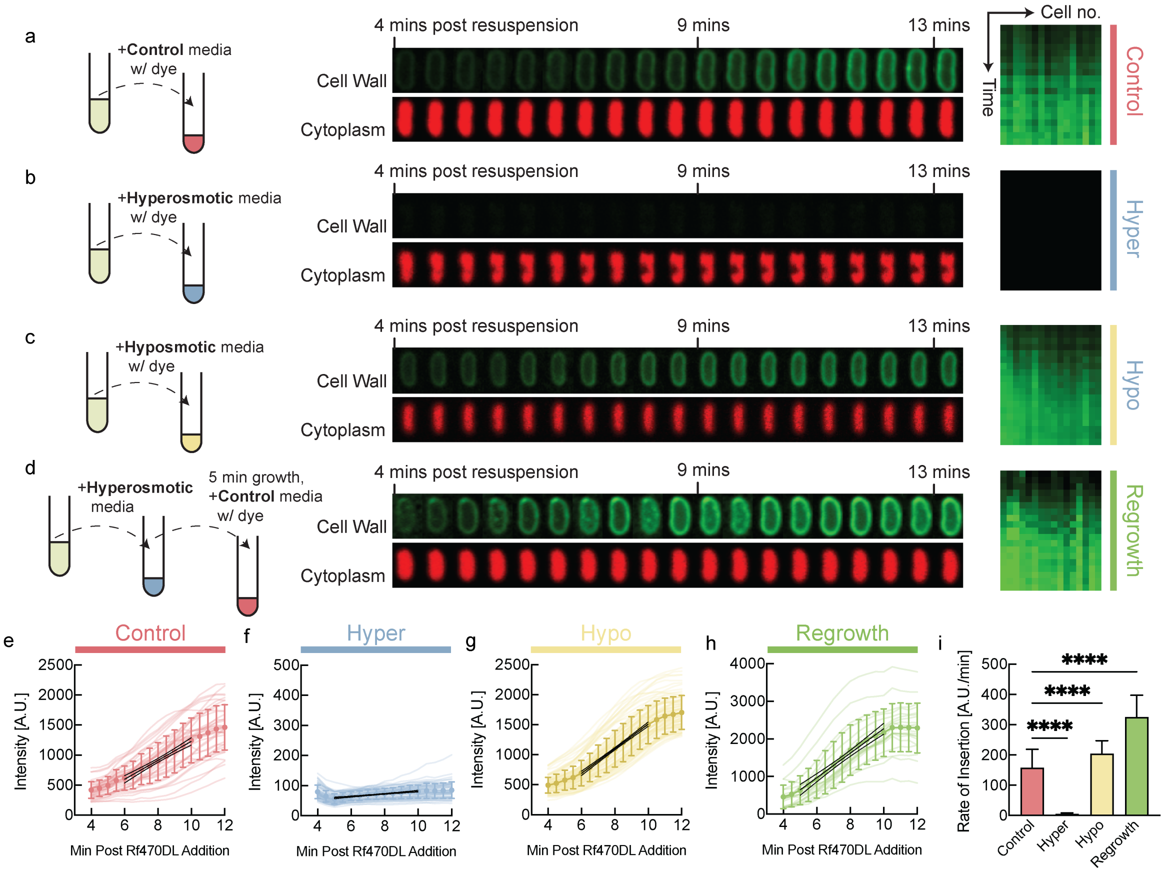

***Fig. S4: Fluorogenic amino acid Rf470DL integration in cell wall under different osmotic condition measured under agarose pad.*** *Rf470DL are D-amino acids that become fluorescent after integration into the cell wall. E. coli cells are grown in minimal media with glucose and shifted to fluorogenic D-amino acid containing media by centrifugation and resuspension in medium containing Rf470DL, as well as different media osmolarity. Resuspended cells were placed under agarose pad and fluorescence images were recorded. Fluorescence intensity per cell wall length was quantified, which denotes the rate of cell wall biosynthesis.* ***a,*** *An example control cell shifted to identical medium with Rf470DL, images were taken after 4 minutes after the resuspension in fluorogenic amino acid containing media. Right, Kymograph of fluorescence intensity of different control cells.* ***b,*** *An example cell, shifted to hyperosmotic medium that induced plasmolysis. Right, kymograph of fluorescence intensity of different bacteria in plasmolysis in hyperosmotic medium. Images showing increase in fluorescence intensity, representing cell wall integration from 4 minutes after resuspension.* ***c,*** *An example cell, shifted to hypoosmotic medium. Right, kymograph of fluorescence intensity of different bacteria.* ***d,*** *An example cell, shifted back to normal osmolarity after being kept in plasmolysis for 5min. Right, Kymograph of fluorescence intensity of different bacteria shifted back to normal medium after 5min in plasmolysis.* ***e,*** *Fluorescence intensity per cell wall length as a function of time for control cells.* ***f,*** *Fluorescence intensity per cell wall length as a function of time for cells in plasmolysis in hyperosmotic medium.* ***g,*** *Fluorescence intensity per cell wall length as a function of time for cells switched to hypoosmotic medium.* ***h,*** *Fluorescence intensity per cell wall length as a function of time for cells shifted back to normal osmolarity after 5min in plasmolysis.* ***l,*** *Rate of cell wall biosynthesis quantified from the slope of the traces in e-h.*

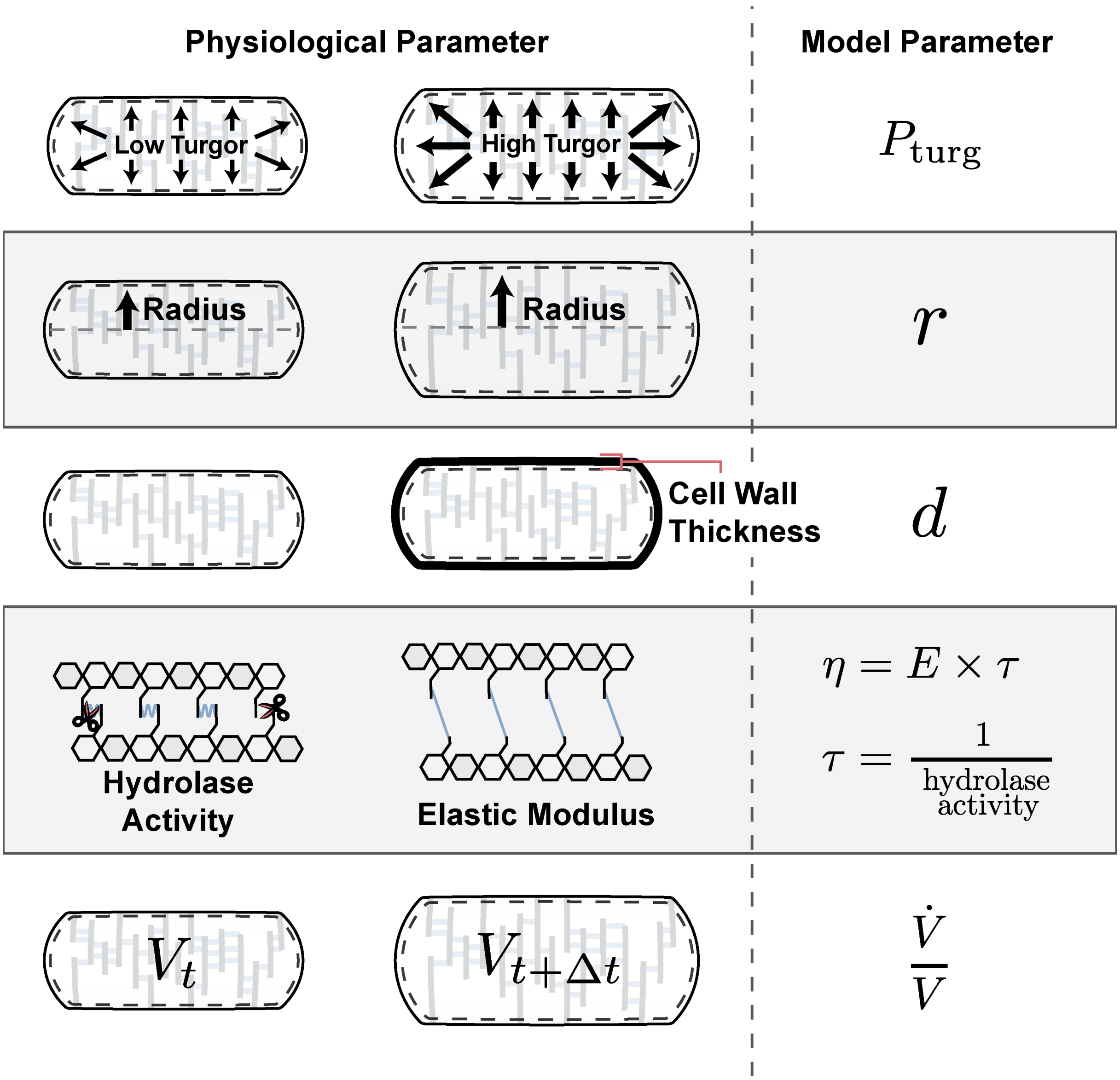

***Fig. S5: Summary of parameters and their physiological interpretation.*** *The model consists of a small number of parameters with specific physiological meanings. Turgor pressure $P_{turg}$ is the osmotic pressure exerted by the cytoplasm on the cell envelope. Cell width is given by two times the radius of the bacterial rod, $r$. The wall thickness $d$ is a dimensional parameter related to the number of peptidoglycan layers. The effective cell wall viscosity $\eta$ is determined by the combination of the elastic modulus E of the peptidoglycan network and the viscoelastic relaxation time $\tau$ that is determined by the remodeling rate of cell wall endopeptidases.*

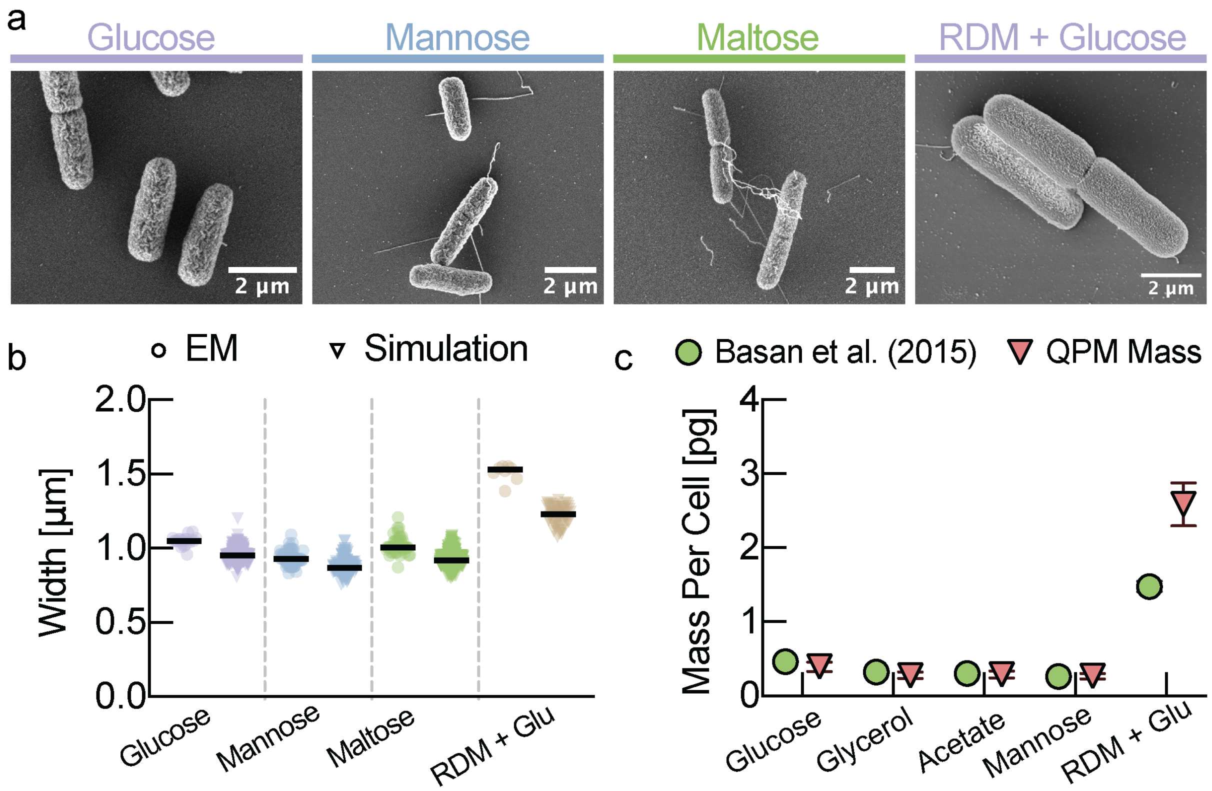

***Fig. S6: Validation of QPM method with traditional experimental methods. a,*** *Electron microscopy (EM) images were taken of E. coli grown in different growth conditions.* ***b,*** *Cell width in different growth conditions quantified from EM images (circles) shows good agreement with widths determined via the QPM simulation method (triangles) (Fig. S3). Individual data points represent individual cells measured.* ***c,*** *Average mass per cell in different growth conditions, measured using traditional methods (circles). Published previously, the data from the traditional method was obtained by baking cell pellets to evaporate water, weighing remaining dry mass and dividing the result by the number of cells determined from both Coulter counting and plating*[25]*. Data from the traditional method shows good agreement with average mass measured via QPM (triangles).*

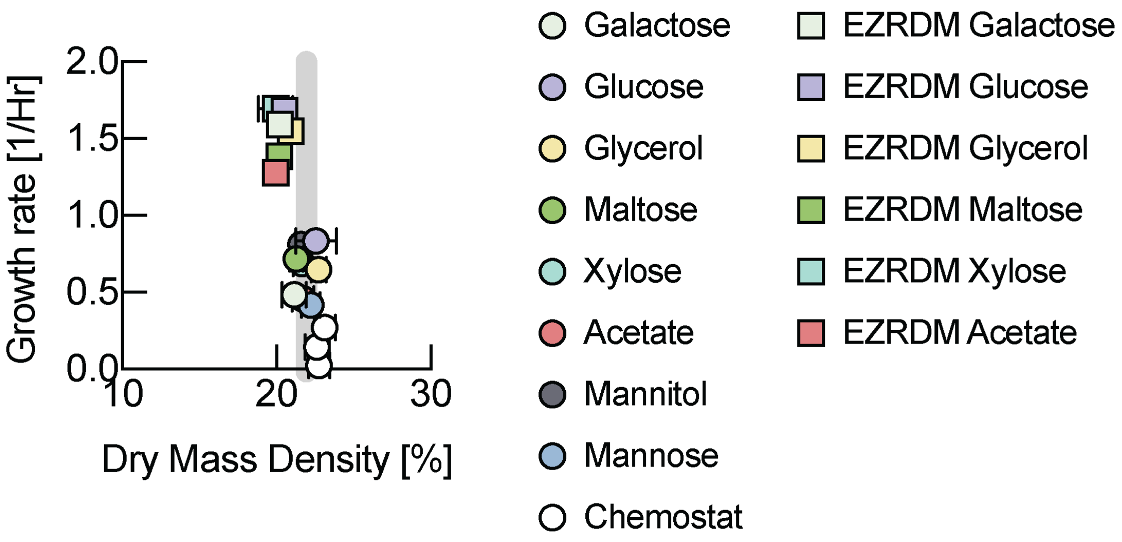

***Fig. S7: Biomass density in different growth conditions including rich defined media.*** *Biomass density is remarkably constant over a wide range of growth rates. We do find a small shift for rich defined media. Error bars in growth rate are the standard deviation from biological replicates (Ez-Acetate: 4 measurements from n=3 biological replicates; Ez-Galactose: 3 measurements from n=2 biological replicates; Ez-Glucose: 3 measurements from n=3 biological replicates; Ez-Glycerol: 3 measurements from n=2 biological replicates; Ez-maltose: 4 measurements from 3 biological replicates; Ez-Xylose: 1 measurement from 1 biological replicate). Replicates for other conditions are summarized in Fig. 3c of the main text. Plotted dry mass density values result from averaging mean dry mass densities for the measured cell populations from different biological replicates. Error bars in dry mass density were determined by propagating the standard deviations of the single cell distributions of the different biological replicates.*

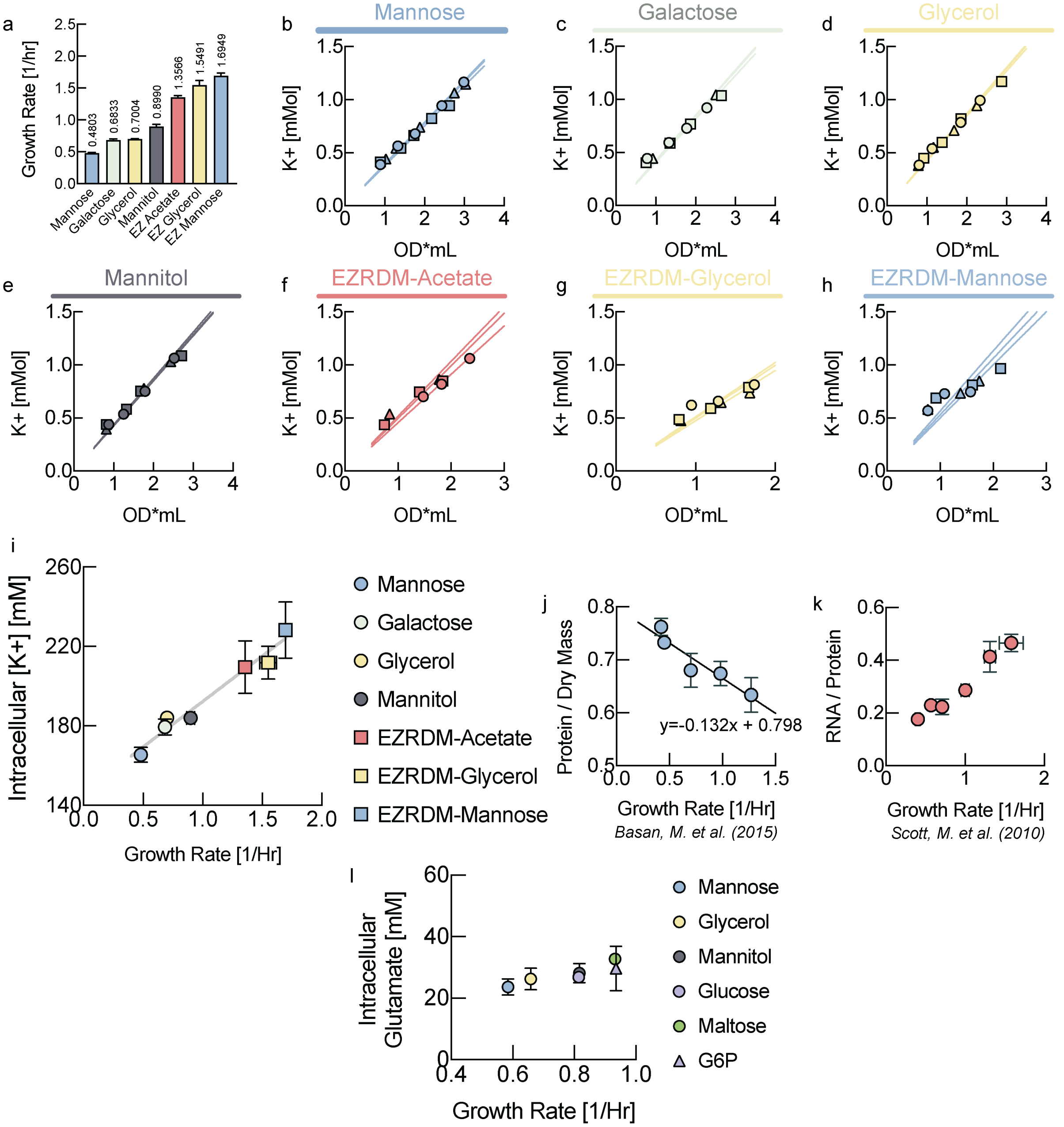

***Fig. S8: Measurements of intracellular potassium. a,*** *Growth rate of E. coli NCM3722 strain grown in low potassium minimal media with different carbon sources. Growth rates were measured from the same samples from which intracellular potassium were measured.* ***b-h,*** *In each growth condition, multiple samples for potassium measurement were taken along the growth curve at different* $OD_{600}$ *from two biological replicates (different symbol shape). The resulting slope of the direct proportionality reflects intracellular potassium concentrations. These measurements allow the comparison of relative concentrations. Because potassium diffuses out of the cell during the wash step, required to remove residual potassium from the medium from the filter, these measurements can only be used to infer relative intracellular concentrations.* ***i,*** *Intracellular potassium concentration as a function of growth rate. Each point represents the mean value of intracellular potassium concentration measured from biological replicates. Error bars S.D.*

***j-k*, Data for conversion to absolute charge concentrations. *j*,** *Replotted data from Basan et al [25]. were used to fit the growth rate dependence of the protein per dry mass.* ***k*,** *Replotted RNA / protein ratios measured by Scott et al. [27] across growth rates.*

***l*,** *Intracellular glutamate concentration as a function of growth rate measured from E. coli NCM3722 strain grown in different carbon sources.*

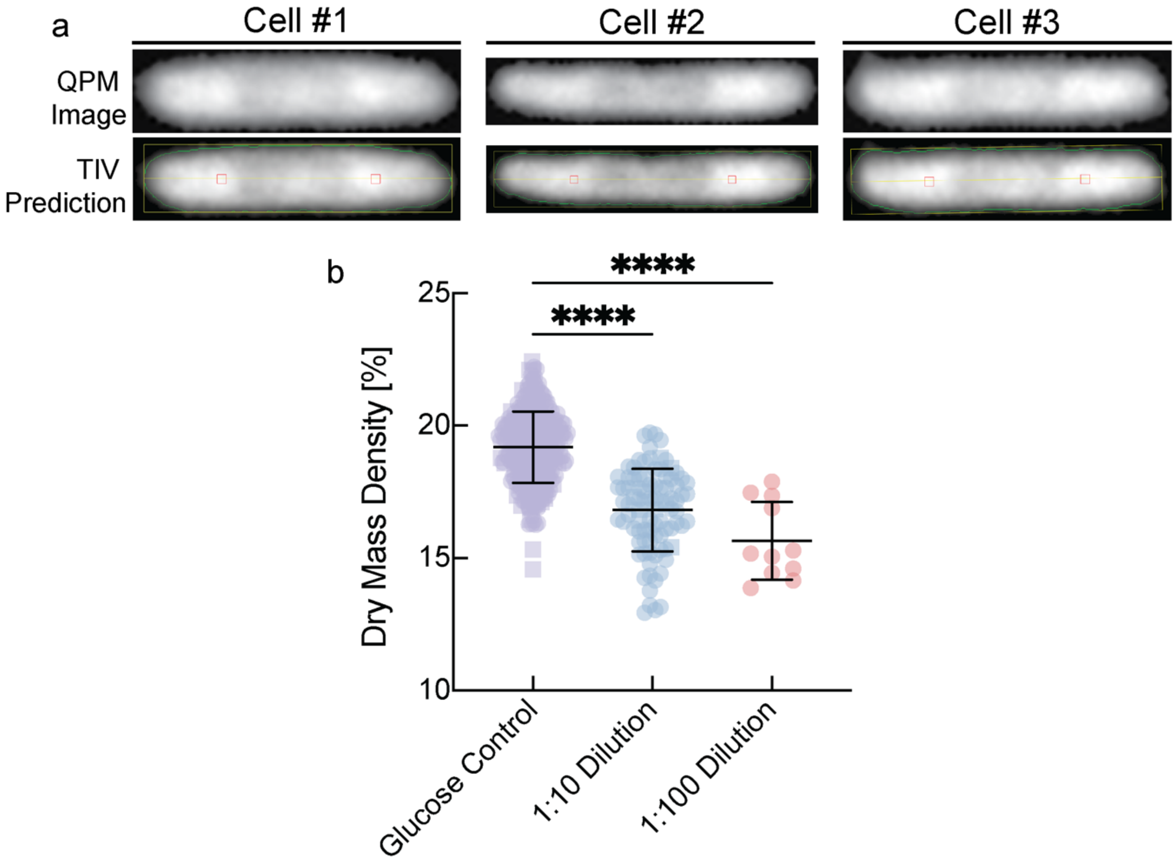

***Fig. S9: TIV method for volume determination of overexpression strains.***

***a,*** *Top panel: QPM images of example bacteria overexpressing large amounts of protein. These bacteria are very long and their shape deviates from a perfect capsule shape. Bottom panel: Cell boundary prediction via TIV method. Predicted cell boundary is marked in thin green line, yellow line denotes the symmetry axis, red squares denote the pixels on symmetry axis which are used for the self-consistent threshold (see supplementary note on TIV prediction.* ***b,*** *Dry mass density of E. coli NCM3722 strain in minimal media with glucose (purple). NCM3722 cells when grown in diluted media (1:10 dilution blue symbols, and 1:100 pink symbols) shows reduced biomass density*.

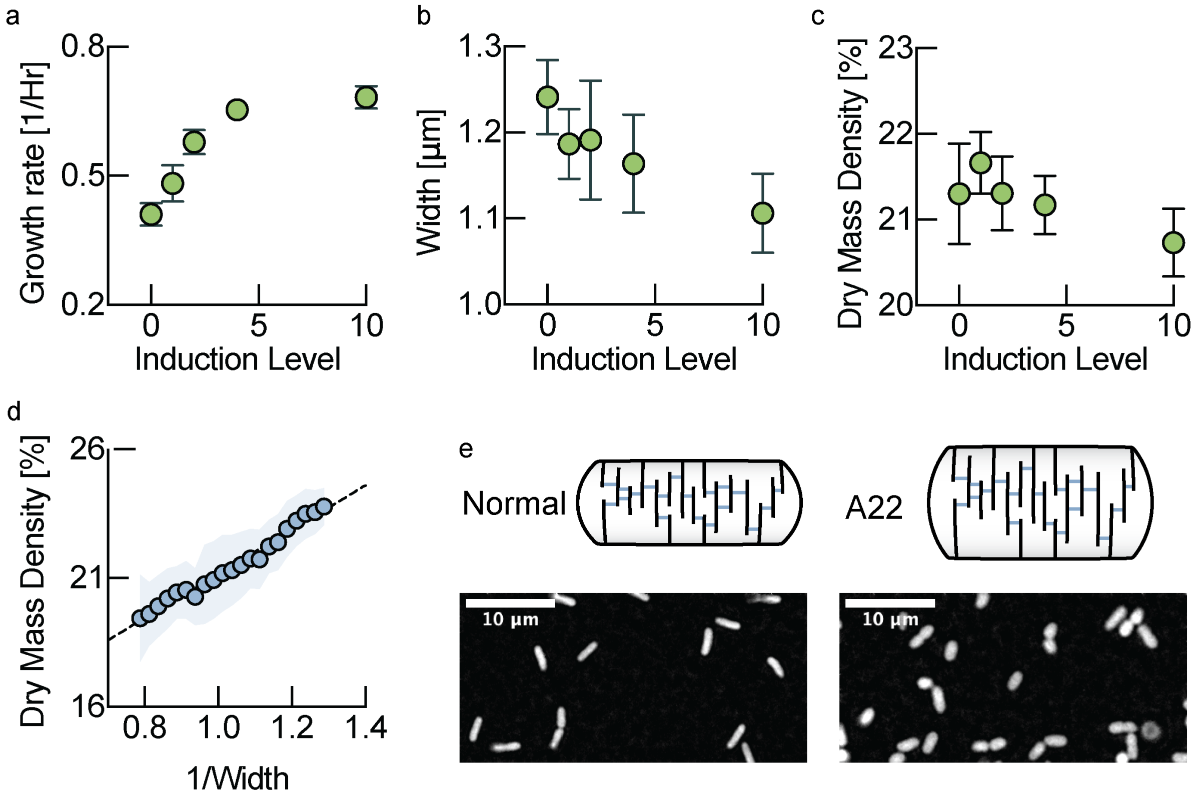

***Fig. S10: Cell wall viscosity can be controlled by endopeptidase titration a,*** *Growth rate as a function of induction levels of endopeptidase expression (number of biological replicates per induction level: 0: 6 replicates; 1: 6 replicates; 2: 5 replicates; 4: 4 replicates; 10: 6 replicates. Error bars represent standard deviation).* ***b,*** *Cell width as a function of induction levels of endopeptidase expression. Biological replicates identical to panel a. Data points are mean of replicates and error bars standard deviations* ***c,*** *Biomass density as a function of induction levels of endopeptidase expression. Biological replicates identical to panel a. Data points are mean of replicates and error bars standard deviations* ***d-e,biomass density dependence on cell width. d,*** *Average biomass density binned for different cell widths for data pooled from wildtype E. coli growing on all different carbon sources. Shaded areas represent the standard deviation of the distribution of cells in each bin. Bin increment: $0.025\mu m^{-1}$.* ***e,*** *another way to perturb cell width is using the antibiotic A22. A22 inhibits MreB, a protein involved in the regulation of cell width. Sub-lethal doses of A22 result in wider cells (right) compared to conditions without A22 (left). Binned data is shown in Fig. 4F of the main text (red triangles).*
